# A lysosomal surveillance response (LySR) that reduces proteotoxicity and extends healthspan

**DOI:** 10.1101/2022.06.13.495962

**Authors:** Terytty Yang Li, Arwen W. Gao, Xiaoxu Li, Yasmine J. Liu, Rachel N. Arey, Kimberly Morales, Amélia Lalou, Qi Wang, Tanes Lima, Johan Auwerx

**Author notes:** These authors contributed equally (Terytty Yang Li, Arwen W. Gao).

## Abstract

Lysosomes are cytoplasmic organelles central for the degradation of macromolecules to maintain cellular homeostasis and health. Here, we discovered an adaptive lysosomal transcriptional response that we termed the Lysosomal Surveillance Response (LySR). Typified by the induction of a large group of transcripts involved in lysosomal function and proteolysis, the LySR can be triggered by silencing of specific vacuolar H^+^-ATPase subunits in *Caenorhabditis elegans*. Notably, LySR activation enhances the clearance of protein aggregates in worm models of Alzheimer’s and Huntington’s disease and amyotrophic lateral sclerosis, thereby boosting fitness and extending lifespan. The GATA transcription factor, ELT-2, regulates the LySR program as well as its associated beneficial effects. In mammalian cells, overexpression of GATA4/GATA6, the mammalian orthologs of ELT-2, is sufficient to induce the expression of multiple lysosome-specific proteases and alleviate proteotoxicity. Activating the LySR pathway may therefore represent an attractive mechanism to reduce proteotoxicity and, as such, potentially extend healthspan.

**Highlights:**

- RNAi of specific v-ATPase subunits extends *C. elegans* lifespan and activates LySR
- GATA transcription factor ELT-2 regulates LySR and LySR-associated lifespan extension
- LySR activation reduces protein aggregates and extends worm healthspan
- Overexpression of GATA4/GATA6 alleviates amyloid-β proteotoxicity in mammalian cells

## INTRODUCTION

Lysosomes are crucial cytoplasmic organelles for degradation and recycling of building blocks, and control multiple cellular signaling and metabolic pathways (Hipp et al., 2019; Lawrence and Zoncu, 2019; Ballabio and Bonifacino, 2020; Saftig and Puertollano, 2021). A variety of substrates are degraded in the lysosomes, ranging from macromolecules including proteins, glycans, lipids and nucleic acids, to organelles and pathogens, which reach the lysosomes either through the endocytic, phagocytic or autophagic routes (Luzio et al., 2007; Mizushima et al., 2008; Wang et al., 2018). The catabolic function of the lysosome is accomplished by a wide repertoire of proteases, lipases, nucleases, sulfatases and other hydrolytic enzymes that usually require an optimal acidic pH of 4.5-5.0, regulating many processes such as the turnover of cellular components, downregulation of surface receptors, inactivation of pathogenic organisms, antigen presentation and bone remodeling.

Dysfunction of lysosomes has been historically associated with lysosomal storage disorders (LSDs), commonly caused by impaired degradation of lysosomal substrates due to mutations in acidic hydrolases as well as non-enzymatic lysosomal proteins (Futerman and van Meer, 2004; Platt et al., 2018; Bonam et al., 2019). Comprising more than 70 individual rare pathologies, the LSDs have a combined incidence of 1 in 5,000 live births and typically manifest progressive neurodegeneration symptoms since infancy or childhood (Nixon et al., 2008; Platt et al., 2018). Accordingly, the accumulation of misfolded and aggregated proteins caused by impaired lysosomal function facilitates the onset and progression of multiple age-related neurodegenerative disorders including Alzheimer disease (AD), Parkinson’s Disease (PD), Huntington’s disease (HD) and Amyotrophic Lateral Sclerosis (ALS) (Balch et al., 2008; Bohnert and Kenyon, 2017; Bonam et al., 2019). Moreover, the aging process itself likely involves a decline in lysosomal activity, and improving lysosomal function may provide promising avenues to extend organismal healthspan (Hughes and Gottschling, 2012; Folick et al., 2015; Carmona-Gutierrez et al., 2016; Leeman et al., 2018; Campisi et al., 2019). For example, in yeast, an early age increase in vacuolar pH limits mitochondrial function and lifespan (Hughes and Gottschling, 2012); in *C. elegans*, the transcription factor EB (TFEB) orthologue HLH-30 regulates lysosomal function, autophagy and longevity (Sardiello et al., 2009; Settembre et al., 2011; Lapierre et al., 2013); while in mouse quiescent neural stem cells and in the germ-cell lineage of *C. elegans*, perturbation of lysosomal activity has been shown to affect the accumulation of protein aggregates and features of stem cell aging (Bohnert and Kenyon, 2017; Leeman et al., 2018).

The vacuolar H^+^-ATPase (v-ATPase), which consists of more than 20 subunits in *C. elegans*, is a highly conserved large complex proton pump at the lysosomal surface essential for the acidification of lysosomes (Forgac, 2007; Lee et al., 2010). The subunits of v-ATPase are generally organized into two domains: a water-soluble ATP-hydrolyzing v1 domain and a membrane-embedded v0 proton channel domain, which function together in coupling the energy of ATP hydrolysis to the transport of protons across the lipid bilayer. The expression of many v-ATPase subunit transcripts decreases with aging (Sun et al., 2020). In addition to its role as a proton pump, v-ATPase has also been shown to be crucial for sensing and integrating of multiple signaling pathways, including the mechanistic target of rapamycin complex 1 (mTORC1) (Zoncu et al., 2011; Lawrence and Zoncu, 2019), adenosine monophosphate-activated protein kinase (AMPK)- metformin (Zhang et al., 2014; Ma et al., 2022) and Janus kinase 2 (JAK2)-signal transducer and activator of transcription-3 (STAT3) signaling (Kreuzaler et al., 2011; Liu et al., 2018), allowing the modulation of key cellular processes such as nutrient sensing, energy metabolism and immune response.

Here, we demonstrate that RNAi of certain lysosomal v-ATPase subunits (e.g., *vha-6*, *vha-8*, *vha- 14*, *vha-20*) extends lifespan by ∼60%, whereas knocking down of some other subunits (e.g., *vha- 1*, *vha-4*, *vha-16* and *vha-19*) shortens lifespan in *C. elegans*. Transcriptomic analysis revealed an up-regulation of 760 genes, enriched for “Lysosome/Proteolysis”, “Metabolic pathways” and “Innate immune response”, specifically in the long-lived *vha-6* RNAi worms. We thereby termed this longevity-linked transcriptional response as the “Lysosomal Surveillance Response (LySR)”. Importantly, *vha-6* RNAi-induced and LySR-associated lifespan extension acts independently of classical longevity regulators, including *daf-2*, *daf-16*, *raga-1*, *aak-2*, *eat-2* and *atfs-1*. Motif prediction analysis of the 760 LySR gene promoters and screening of 14 known GATA transcription factors identified ELT-2 as the key regulator of the LySR and LySR-linked longevity. Moreover, in models of neurodegenerative diseases and of normal aging, *vha-6* RNAi-mediated LySR activation enhances proteostasis, reduces protein aggregates and improves animal health. Finally, we demonstrate that a similar beneficial transcriptional response is also induced by overexpression of GATA4/GATA6 in mammalian cells, leading to increased expression of a set of lysosomal proteases and reduced amyloid-β proteotoxicity. Collectively, these findings provide evidence for a previously unknown mechanism to boost lysosomal function, reduce proteotoxicity, and potentially benefit organismal health against neurodegenerative diseases and normal aging.

## RESULTS

### Knockdown of specific v-ATPase subunits extends *C. elegans* lifespan and activates LySR

In light of the fact that an adaptative anti-aging mitochondrial stress response is activated by RNAi-mediated silencing of *cco-1*, a gene that encodes a mitochondrial respiratory chain complex IV subunit (Dillin et al., 2002; Durieux et al., 2011), while the ER stress response is induced by RNAi of an ER chaperone gene, *hsp-3* (Kapulkin et al., 2005), we asked whether a lysosomal adaptive transcriptional response could be activated by knocking down specific v-ATPase subunits and explored its potential implications in health and longevity. By measuring the lifespan of *C. elegans* fed with RNAi targeting each of the major v-ATPase subunits (Figures 1A-1G, and S1A-S1M), we found that *vha-6* RNAi extended lifespan by almost 80%, while less pronounced lifespan extensions were detected in worms fed with *vha-8*, *vha-14*, *vha-15* or *vha-20* RNAi (Figures 1A- 1E). On the contrary, RNAi targeting *vha-1*, *vha-4*, *vha-16* or *vha-19* shortened lifespan (Figures 1F-1G, S1A and S1D).

**Figure 1.**
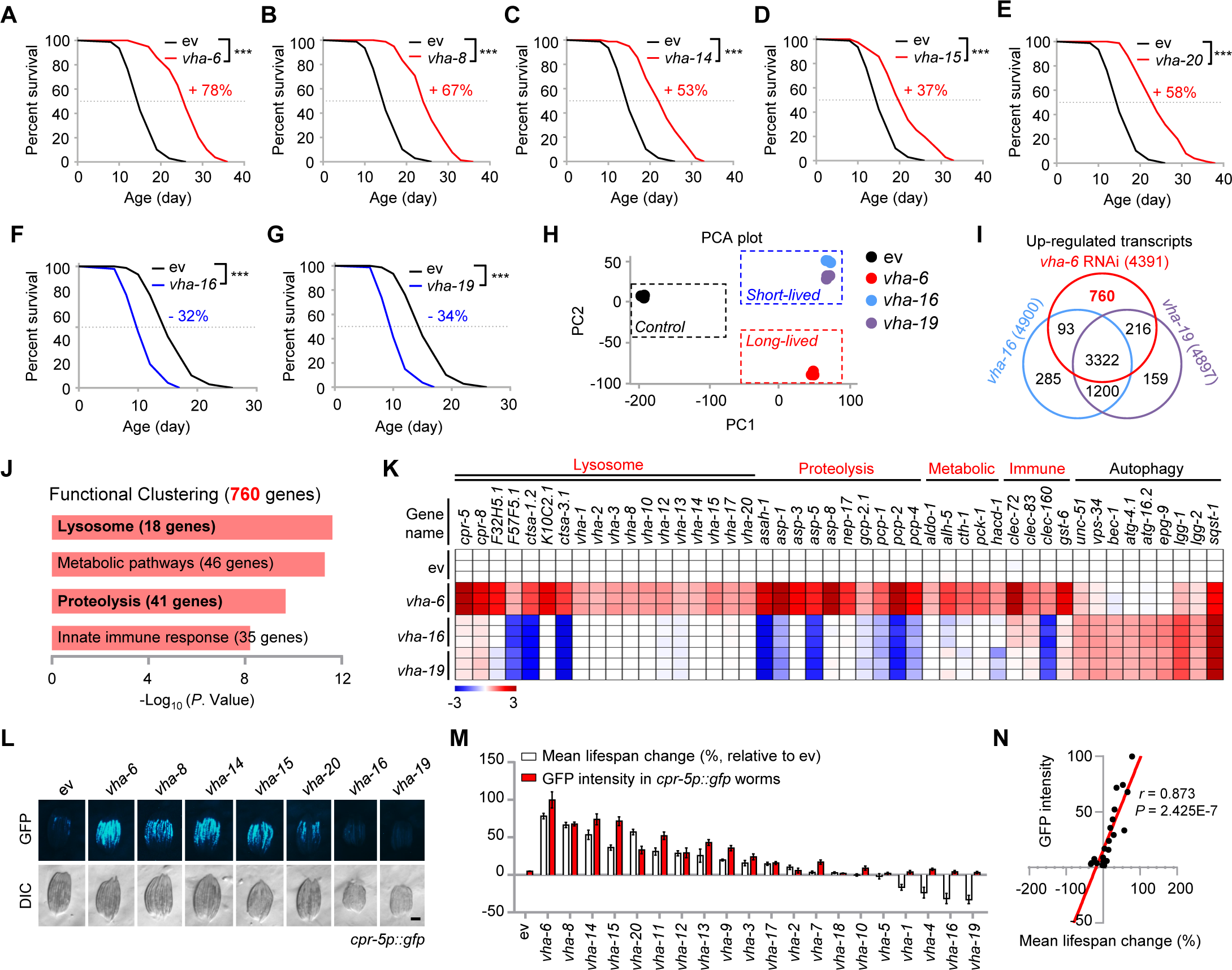
Knockdown of specific v-ATPase subunits extends *C. elegans* lifespan and activates LySR. (**A** to **G**) Survival of worms fed with control (ev), or RNAi targeting *vha-6* (A), *vha-8* (B), *vha-14* (C), *vha-15* (D), *vha-20* (E), *vha-16* (F) and *vha-19* (G). Each v-ATPase RNAi occupied 40%, except for *vha-6* (20%), *vha-16* (10%) and *vha-20* (20%) RNAi. Control RNAi was used to supply to a final 100% of RNAi for all conditions. The percentages indicate the mean lifespan changes relative to control. (**H**) Principal-component analysis (PCA) plot of the RNA-seq results of the RNA-seq results of the worms fed with control, *vha-6* (long-lived), *vha-16* and *vha-19* (short-lived) RNAi. (**I**) Venn diagram of the up-regulated differentially expressed genes (DEGs) in response to *vha-6*, *vha-16* and *vha-19* RNAi. (**J**) Functional clustering of the 760 DEGs as indicated in (I). (**K**) Heat-map of the relative expression levels of representative DEGs in response to *vha-6*, *vha-16* and *vha-19* RNAi. The color represents gene expression differences in log_2_(fold change, FC) relatively to the control RNAi condition. (**L**) GFP expression levels of *cpr-5p::gfp* worms fed with RNAi targeting different v-ATPase subunits. Scale bar, 0.3 mm. (**M**) Percentages of the mean lifespan change (relative to ev condition) and GFP intensity of *cpr-5p::gfp* worms fed with control, or RNAi targeting v-ATPase subunits (*n* = 3 independent experiments). (**N**) GFP intensity of *cpr-5p::gfp* worms positively correlates with worm lifespan change. Pearson’s correlation coefficient (*r*) was calculated with the mean lifespan change values (*x* axis) and the GFP intensity of *cpr-5p::gfp* worms (*y* axis) as indicated in (M). Error bars denote SEM. Statistical analysis was performed by log-rank test in (A to G) (\*\*\**P* < 0.001). See also Figure S1 and Table S1.

To determine the footprints underlying the lifespan changes conferred by silencing different v-ATPase subunits, we compared the transcript profiles of *C. elegans* fed with *vha-6* (extended lifespan), and *vha-16* or *vha-19* (shortened lifespan) RNAi (Table S1). Principal-component analysis (PCA) of the RNA-seq results revealed that worms with different lifespan indeed had separated gene expression signatures (Figure 1H). More specifically, knockdown of each of the three v-ATPase subunits induced the up-regulation (Log2FC > 1, adjusted *P* < 0.05) of 4,391-4,900 genes, and the majority (3,322 genes) of them were shared (Figure 1I). Gene ontology (GO) analysis revealed that 898 (27.0%) of the 3,322 genes were related to “Integral component of membrane” (Figure S1N), confirming a common and key role of the v-ATPase subunits in maintaining the v-ATPase-mediated membrane-associated biological processes (Forgac, 2007). Similarly, 3,026 genes were commonly down-regulated (Log2FC < -1, adjusted *P* < 0.05) upon the three v-ATPase RNAi and were overall related to “Nucleus” and “Embryo development” (Figures S1O and S1P). In particular, 760 genes were exclusively up-regulated in the long-lived *vha-6* RNAi model, but not in the short-lived *vha-16* or *vha-19* RNAi model (Figure 1I). These 760 genes were enriched for “Lysosome/Proteolysis”, “Metabolic pathways” and “Innate immune response” (Figures 1J and 1K). We named this unique longevity-linked transcriptional response, the “<u>Ly</u>sosomal <u>S</u>urveillance <u>R</u>esponse (LySR)”, which can be triggered by knocking down specific v-ATPase subunits (e.g., *vha-6*), and is typified by the strong induction of a large panel of genes related to lysosome and proteolysis, such as *cpr-5* and *cpr-8,* two worm orthologs of human Cathepsin B (CTSB) (Yadati et al., 2020). Of note, the LySR program covered a variety of endopeptidase types including the cysteine-type (e.g., *cpr-5*), serine-type (e.g., *ctsa-1.2*), aspartic-type (e.g., *asp-1*), metallo-type (e.g., *nep-17*), dipeptidyl-type (e.g., *pcp-1*) as well as amidohydrolase (e.g., *asah-1*) (Figure 1K). Interestingly, while RNAi of *vha-6*, *vha-16* and *vha- 19* all induced the expression of some autophagy-related transcripts, *vha-16* or *vha-19* RNAi worms had even higher levels of autophagy genes as compared to that in *vha-6* RNAi worms (Figure 1K), confirming different metabolic profiles in the long-lived and short-lived worms.

Notably, the GFP intensity of *cpr-5p:gfp* worms fed with RNAi against different v-ATPase subunits strongly correlated with the changes in their mean lifespans (Pearson’s *r*, *P* = 2.425 × 10^−7^) (Figures 1L-1N, and S1Q), indicating that the transcriptional level of the lysosomal protease, CPR-5, is likely predictive for the longevity of v-ATPase RNAi worms.

As expected, the expression of lysosome/proteolysis related transcripts, including *cpr-5*, *cpr-8*, *ctsa-1* and *asp-10*, robustly increased in the long-lived (e.g., *vha-6, vha-8, vha-14, vha-15* and *vha- 20* RNAi) worms, but only minorly affected in the short-lived (e.g., *vha-16* and *vha-19* RNAi) worms (Figure 2A). The impact of *vha-6* RNAi on GFP induction in *cpr-5p:gfp* worms and lifespan extension was reliably reproduced when worms were exposed to different amounts of *vha- 6* RNAi (Figures 2B and 2C). Moreover, another two RNAi clones (*vha-6*_RNAi_2 and *vha- 6*_RNAi_3) targeting different regions of the *vha-6* mRNA, as compared to that used in the RNAi screening (*vha-6*_RNAi_1) (Figure 2D), consistently induced GFP-CPR-5 expression and extended the lifespan (Figures 2E and 2F). Furthermore, *vha-6* RNAi extended worm lifespan even when the RNAi treatment started since the L4/young adult stage (Figure S1R).

**Figure 2.**
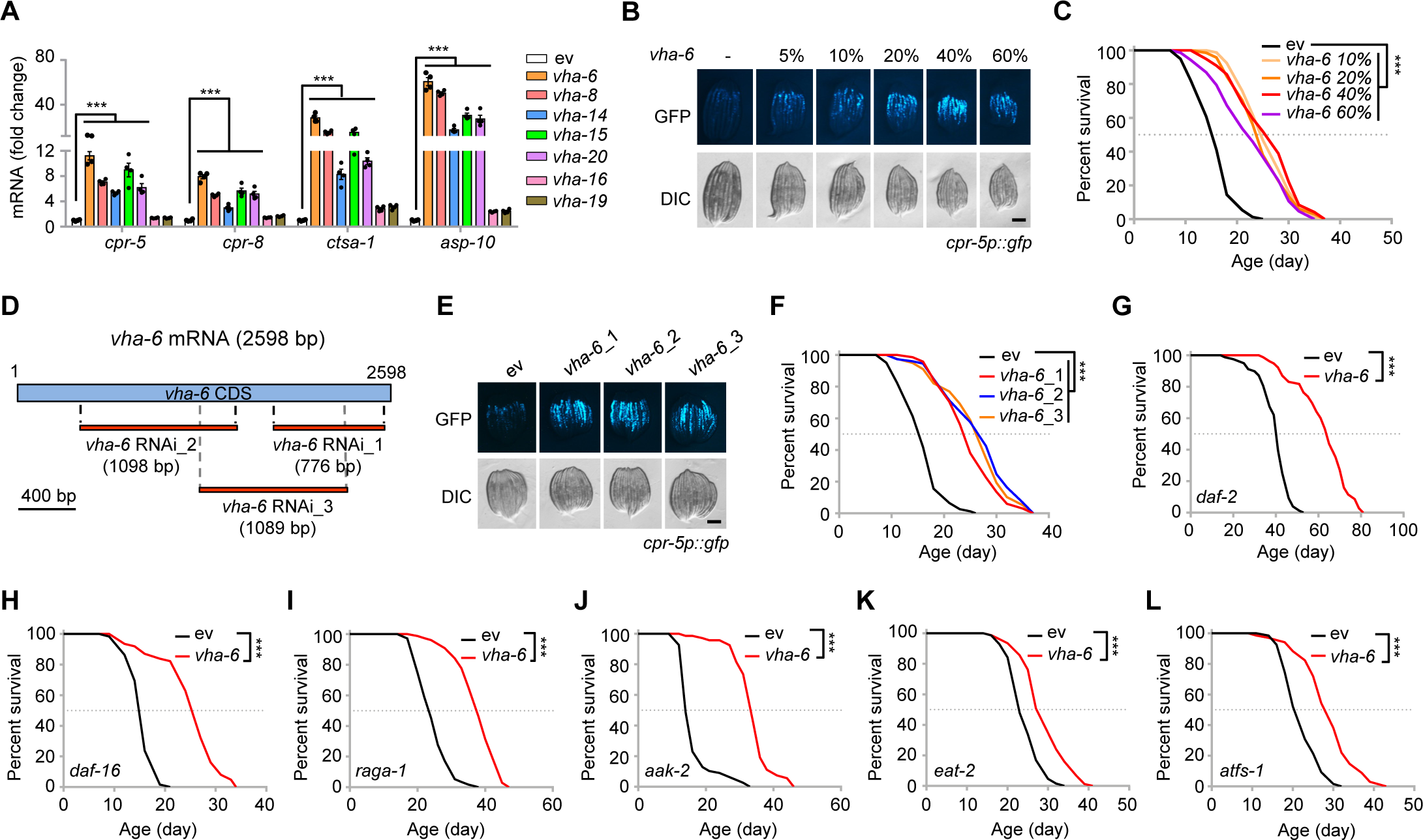
Impact of *vha-6*, *-8*, *-14*, *-15*, *-20*, *-16* and *-19* RNAi on gene expression and lifespan of *C. elegans*. (**A**) qRT-PCR analysis of transcripts (*n* = 4 biologically independent samples) in worms fed with control (ev), or RNAi targeting v-ATPase subunits. (**B** and **C**) GFP-CPR-5 expression level (B) and survival (C) of worms fed with control, or 10%-60% *vha-6* RNAi, control RNAi was used to supply to a final 100% of RNAi for all conditions. Scale bar, 0.3 mm. (**D**) Schematic diagram showing the regions on mRNA targeted by the three *vha-6* RNAi obtained from either the Vidal (*vha-6*_1) or Ahringer (*vha-6*_2, *vha-6*_3) library. (**E** and **F**) GFP-CPR-5 expression level (E) and survival (F) of worms fed with control or the *vha-6* (20%) RNAi as indicated in (D). Scale bar, 0.3 mm. (**G** to **L**) *vha-6* RNAi extends the lifespan of *daf-2(e1370)* (G), *daf-16(mu86)* (H), *raga-1(ok386)* (I), *aak-2(ok524)* (J), *eat-2(ad465)* (K) and *atfs- 1(gk3094)* (L) mutants by 61%, 68%, 63%, 138%, 20% and 38%, respectively. Error bars denote SEM. Statistical analysis was performed by ANOVA followed by Tukey post-hoc test in (A), or by log-rank test in (C), (F to L) (\*\*\**P* < 0.001).

### RNAi of *vha-6* extends worm lifespan independent of classical longevity pathways

To test if any of the canonical longevity pathways contribute to *vha-6* RNAi-induced lifespan extension, we knocked down *vha-6* in worms carrying null mutations in insulin/IGF-1 signaling (Kenyon et al., 1993), mTOR signaling (Schreiber et al., 2010), AMPK signaling (Schulz et al., 2007), caloric restriction (Lakowski and Hekimi, 1998) and mitochondrial stress signaling (Nargund et al., 2012). *vha-6* RNAi extended the lifespan of *daf-2*, *daf-16*, *raga-1*, *aak-2, eat-2* and *atfs-1* mutants (Figures 2G-2L), suggesting that *vha-6* RNAi regulates longevity independently of insulin/IGF-1 (*daf-16/daf-2*), mTOR/AMPK signaling (*raga-1*/*aak-2*), caloric restriction (*eat-2*) and mitochondrial stress response (*atfs-1*) pathways. Strikingly, the mean lifespan of *vha-6* RNAi fed *daf-2(e1370)* was extended to 64 days (Figure 2G), 1.6-fold and 3.6- fold greater than that of the control RNAi fed *daf-2(e1370)* (40 days) and wild-type (18 days) worms, respectively.

### Identification of the GATA transcription factor ELT-2 as a key regulator for LySR

To identify which transcription factor governs the LySR activation, we analyzed the promoters of the 760 genes up-regulated only upon *vha-6* RNAi, but not upon *vha-16* or *vha-19* RNAi, and identified a 10-base pair (bp) ACTGATAAGA motif highly enriched in this set of promoters (253 genes out of 760; *P* = 1 × 10^−31^) (Figure 3A). This motif was present as either a single copy or as multiple copies (Table S1), and was located ∼100 bp upstream of the transcription start sites (TSS) for both the 760 “*vha-6* only” genes and all the other *C. elegans* genes, with the “*vha-6* only” genes more enriched according to the similarity scores calculated based on the position weight matrix (PWM) (Figure 3B). We next asked which transcription factors may bind to this motif. After comparing this motif with the putative binding motifs of all known transcription factors in *C. elegans*, 8 GATA transcription factors were found among the top 10 hits (Figure 3C), in line with the presence of a “GATA” sequence at the center of the 10-bp motif (Figure 3A).

**Figure 3.**
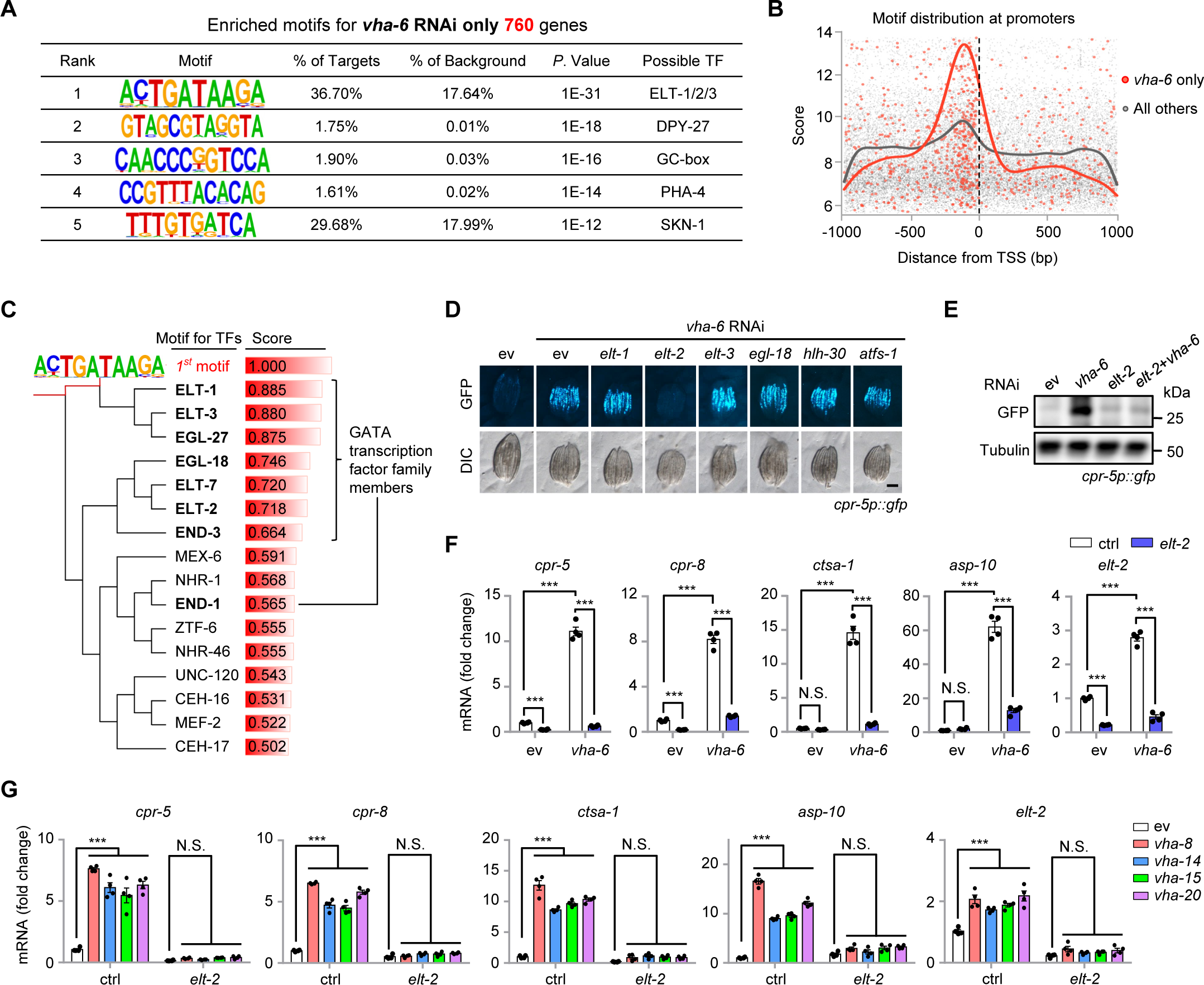
Identification of the GATA transcription factor, ELT-2, as a key regulator for LySR. (**A**) The top five enriched motifs for promoters of the 760 up-regulated genes upon *vha-6* RNAi, but not upon *vha-16* or *vha-19* RNAi, by using Hypergeometric optimization of motif enrichment (HOMER) and ranked based on the *P* values. (**B**) Genomic distribution of the motif hits at the promoters of the *vha-6* RNAi only genes (red), and all other *C. elegans* genes (gray). Scores are assigned based on the position weight matrix (PWM) of the motif as shown in (A). Lines indicate the density of the motif hits. TSS, transcription start site. (**C**) The similarity of the Rank 1 motif as found in (A) to the putative binding motifs of all known transcription factors in *C. elegans*. All transcription factors with similarity score above 0.500 were shown. The GATA transcription factor family members were highlighted in bold. (**D**) RNAi of *elt-2* attenuated the GFP expression of *cpr-5p::gfp* worms induced by *vha-6* RNAi. RNAi targeting *vha-6* occupied 40%, RNAi targeting *elt-1*, *elt-2*, *elt-3*, *egl-18*, *hlh-30* and *atfs-1* occupied 60%. Scale bar, 0.3 mm. (**E** and **F**) Western blots (E) and qRT-PCR analysis of transcripts (*n* = 4 biologically independent samples) (F) in *cpr-5p::gfp* worms fed with control or *vha-6* RNAi, and/or *elt-2* RNAi. (**G**) qRT-PCR analysis of transcripts (*n* = 4 biologically independent samples) in worms fed with control, *vha-8*, *vha-14*, *vha-15* and *vha-20* RNAi, and/or *elt-2* RNAi. Error bars denote SEM. Statistical analysis was performed by ANOVA followed by Tukey post-hoc test (\*\*\**P* < 0.001; N.S., not significant). See also Figure S2 and Table S1.

By feeding worms with RNAi’s targeting all 14 known GATA transcription factor genes in *C. elegans* (Budovskaya et al., 2008; Mann et al., 2016), we discovered that *elt-2* RNAi, but not others, almost completely blocked the GFP induction in *cpr-5p:gfp* worms in response to *vha-6* silencing (Figures 3D, 3E, and S2A). In addition, RNAi of the essential mitochondrial unfolded protein response (UPR^mt^) transcription factor *atfs-1* (Nargund et al., 2012); *hlh-30,* the worm ortholog of the key lysosomal gene regulator TFEB (Sardiello et al., 2009; Lapierre et al., 2013); or *pqm-1*, which encodes a transcription factor that binds to a GATA-like DAF-16 Associated Element (DAE) motif (Tepper et al., 2013), did not affect *vha-6* RNAi-induced GFP expression in *cpr-5p:gfp* worms (Figures 3D and S2A). Knockdown of *elt-2* furthermore abrogated the induction of lysosome/proteolysis-related transcripts, including *cpr-5*, *cpr-8*, *ctsa-1* and *asp-10*, upon RNAi of *vha-6*, *vha-8*, *vha-14*, *vha-15* or *vha-20* individually (Figures 3F and 3G). In contrast, the *vha-6* RNAi-induced expression of these lysosomal proteases was not affected in autophagy-defective mutants (Figure S2B).

### ELT-2 determines LySR activation and LySR-associated lifespan extension

We then performed another RNA-seq experiment, this time using the alternative *vha-6* RNAi_2, which targets a different and broader region of the *vha-6* mRNA as compared to the *vha-6* RNAi_1 previously used in Figure 1I (Figure 2D). 3,201 transcripts were commonly induced by *vha-6* RNAi_1 and *vha-6* RNAi_2, among which 488 genes overlapped with the 760 LySR genes up- regulated with *vha-6* RNAi, but not with *vha-16* or *vha-19* RNAi (Figure 4A and Table S2). Importantly, within these 488 transcripts, up to 65.2% (318) transcripts required ELT-2 for induction (*P* < 2.2 × 10^−16^; Fisher’s exact test), while only 21.9% (1,229) transcripts rely on ELT- 2 among all the 5,617 *vha-6* RNAi_2-induced transcripts in general (Figure 4A). These ELT-2- dependent transcripts were highly enriched for “Lysosome/Proteolysis”, “Innate immune response” (Shapira et al., 2006; Block et al., 2015), and “Metabolic pathways” (Figures 4B and 4C), in line with the LySR-linked terms (Figures 1J and 1K).

**Figure 4.**
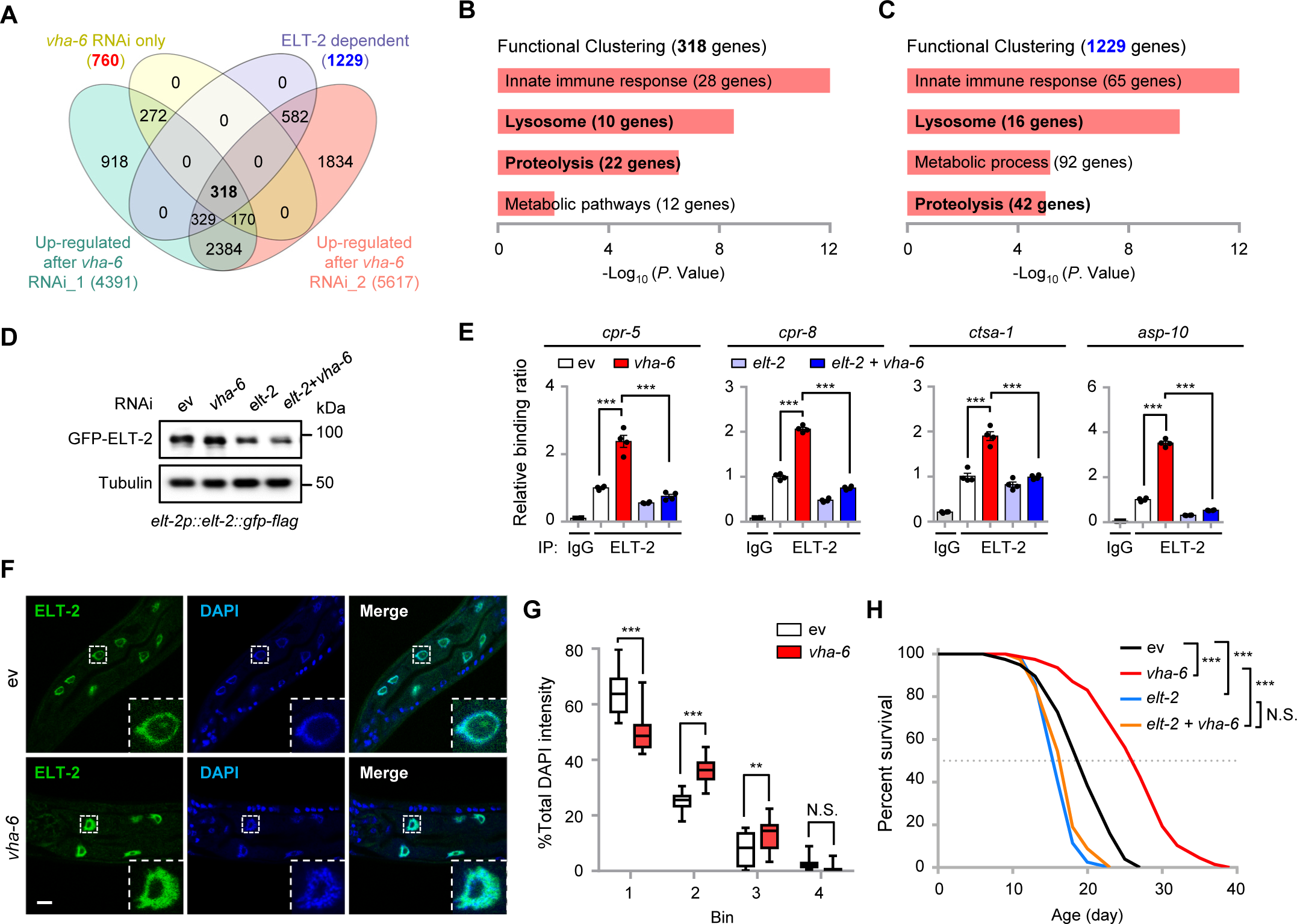
ELT-2 determines LySR activation and LySR-associated lifespan extension. (**A**) Venn diagram of the up-regulated DEGs in response to *vha-6* RNAi_1 or *vha-6* RNAi_2, in common with the ELT-2-dependent DEGs upon *vha-6* RNAi_2, and the 760 “*vha-6* RNAi only” DEGs. (**B** and **C**) Functional clustering of the 318 (B) and 1,229 (C) DEGs as indicated in (A). (**D**) Western blots of *elt-2p::elt-2::gfp-flag* worms fed with control (ev) or *vha-6* RNAi, and/or *elt-2* RNAi. (**E**) ChIP-qPCR (*n* = 4 biologically independent samples) of *elt-2p::elt-2::gfp-flag* worms fed with control (ev) or *vha-6* RNAi, and/or *elt-2* RNAi. ChIP was performed by using IgG control or anti-Flag M2 beads. (**F**) Representative images of GFP-tagged ELT-2 and DAPI staining in *elt-2p::elt-2::gfp-flag* worms fed with control (ev) or *vha-6* RNAi. Scale bar, 10 μm. (**G**) Quantification of the distribution of DAPI or ELT-2 signal in intestinal nuclei in *elt-2p::elt-2::gfp-flag* worms fed on control (ev) or *vha-6* RNAi. Image voxels were ranked by DAPI intensity within each nucleus and divided into 4 equal-volume bins. Percentage of total DAPI intensity in each of the bins was measured (*n* = 30 nuclei for each condition). (**H**) Survival of worms fed with control (ev), *vha-6* RNAi, and/or *elt-2* RNAi. Error bars denote SEM. Statistical analysis was performed by ANOVA followed by Tukey post-hoc test in (E) and (G), or by log-rank test in (H) (\*\**P* < 0.01; \*\*\**P* < 0.001; N.S., not significant). See also Table S2.

Despite a ∼3-fold increase in *elt-2* mRNA level in response to *vha-6* RNAi (Figure 3F), the protein level of ELT-2::GFP was largely unaffected by *vha-6* RNAi in *elt-2p::elt-2::gfp-flag* worms (Figure 4D). In contrast, by using chromatin immunoprecipitation coupled with quantitative PCR (ChIP-qPCR) analysis, robust enrichments of ELT-2 were detected at the promoters of lysosomal proteases including *cpr-5*, *cpr-8*, *ctsa-1* and *asp-10* in response to *vha-6* silencing (Figure 4E), suggesting that ELT-2 directly binds to the promoters of these LySR genes upon *vha-6* RNAi. In line with these observations, we found that although the intestinal ELT-2::GFP strongly co- localized with the blue-fluorescent DNA stain DAPI under both basal and *vha-6* RNAi conditions, the DAPI signal appeared to be more scattered in response to *vha-6* RNAi (Figure 4F). Quantification of the four fractions, divided based on the DAPI staining intensity within each of the nucleus, confirmed an overall higher dispersal and likely less-compacted chromatin/DNA distribution in *vha-6* RNAi-treated intestinal nuclei (Figure 4G). These results suggest some reorganizations of the chromatin/DNA to facilitate the binding of ELT-2 to the promoters of LySR genes upon *vha-6* RNAi, reminiscent of the chromatin changes during mitochondrial perturbations (Merkwirth et al., 2016; Tian et al., 2016).

Importantly, RNAi of *elt-2* almost completely abolished the lifespan extension induced by *vha-6* RNAi (Figure 4H). In line with previous results (Zhang et al., 2013; Mann et al., 2016), *elt-2* RNAi alone shortened lifespan (Figure 4H), which may be due to the decreased basal expression of some of the LySR genes, including *cpr-5* and *cpr-8* (Figures 3F and 3G) (Mann et al., 2016). Overexpression (OE) of *elt-2* with its own promoter has also been shown to extend worm lifespan by ∼20% (Zhang et al., 2013; Mann et al., 2016). However, such extent of lifespan extension is much less than that of the effect of *vha-6*, *vha-8*, *vha-14* or *vha-20* RNAi, which is ∼60% extension (Figures 1A-1E). Consistently, by re-analyzing an extant RNA-seq dataset (GSE69263) (Mann et al., 2016), we found that the LySR genes are only sporadically induced in both young and aged ELT-2 OE worms as compared to that in control worms (Figure S2C). Thus, the GATA transcription factor ELT-2 is an essential, though apparently not the only, regulator of LySR and LySR-associated longevity in *C. elegans*.

### LySR activation reduces protein aggregates and extends the healthspan of *C. elegans*

Lysosomal proteases, including the cathepsins (Yadati et al., 2020), are central enzymes involved in the proteolytic degradation of misfolded and aggregation-prone proteins, such as amyloid-β (Aβ) and polyglutamine (polyQ)-expanded huntingtin (HTT), the contributing factors in the pathogenesis of Alzheimer’s disease (AD) and Huntington’s disease (HD) (Cataldo and Nixon, 1990; Saido and Leissring, 2012), respectively. We thus questioned whether LySR activation could reduce proteotoxicity *in vivo*.

The GMC101 strain is a worm AD model which expresses the human Aβ1-42 peptide in the body-wall muscle cells (McColl et al., 2012). GMC101 adults develop age-progressive paralysis and exacerbated amyloid deposition after a temperature shift from 20 to 25 °C. In response to the temperature shift, transcript levels of multiple LySR-associated lysosomal proteases increased in GMC101 worms (Figure 5A), suggesting that the LySR is induced concomitantly with proteotoxicity. RNAi of *vha-6* further increased the mRNA level of these proteases by more than 10-fold (Figure 5B), and reduced Aβ aggregates in GMC101 worms to an almost undetectable level (close to that in the non-Aβ expressing CL2122 control worms, with both *vha-6_1* and *_2* RNAi) at both 20 and 25°C (Figure S3A), an effect that was abrogated by *elt-2* RNAi (Figure 5C).

**Figure 5.**
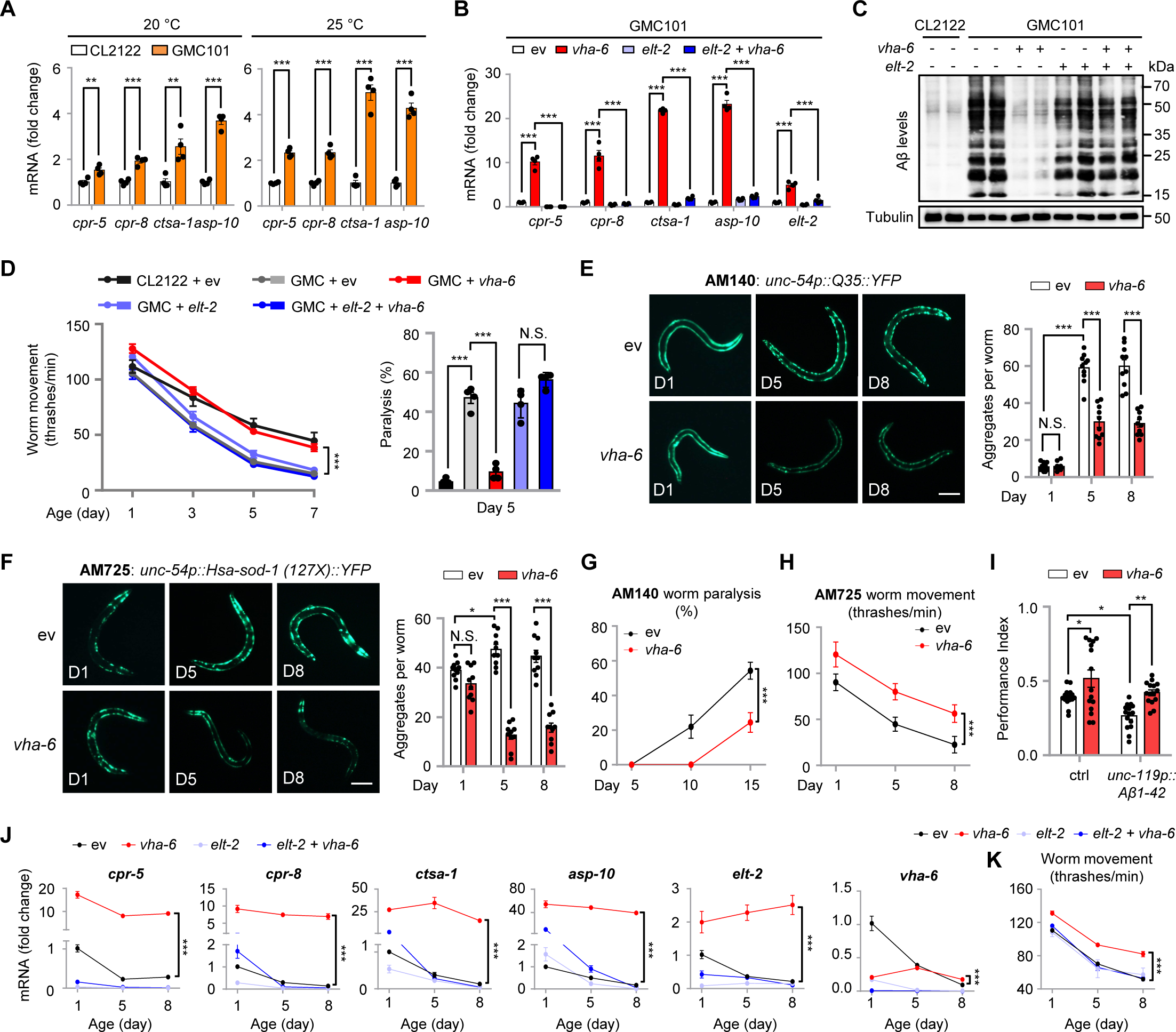
Activation of LySR reduces protein aggregates and extends the healthspan of *C. elegans*. (**A**) qRT-PCR analysis of transcripts (*n* = 4 biologically independent samples) of CL2122 and GMC101 worms cultured at two different temperatures since Larval 4 (L4) stage. (**B** to **D**) qRT-PCR analysis of transcripts (*n* = 4 biologically independent samples) (B), western blots of amyloid-β (6E10) (C), movement (*n* = 12 individual worms for each condition) and paralysis (*n* = 4 independent experiments) (D) of CL2122 or GMC101 worms fed with control, *vha-6* and/or *elt-2* RNAi. (**E** and **F**) RNAi of *vha-6* (20%) reduces the aggregate formation in *unc-54p::Q35::YFP* (polyQ model) (E) and *unc-54p::Hsa-sod-1::YFP* (amyotrophic lateral sclerosis (ALS) model) (F) worms (*n* = 10 individual worms for each condition). Scale bars, 0.2 mm. (**G** and **H**) Paralysis (*n* = 10 independent worm plates for each condition) (G), or movement (*n* = 15 individual worms for each condition) (H) of worms as indicated in (E) and (F). (**I**) *vha-6* RNAi improves intermediate-term memory in the worm AD model GRU102 (*unc-119p::Aβ1-42*) strain with constitutive neuronal Aβ1-42 expression, analyzed at Day 4 adulthood (*n* = 15 chemotaxis assays of 50-100 worms for each condition). (**J** and **K**) mRNA levels of indicated genes (*n* = 4 biologically independent samples) (J), and movement (*n* = 12 individual worms for each condition) (K) of worms fed control or *vha-6* RNAi, in combination with *elt-2* RNAi, harvested at different ages. Error bars denote SEM. Statistical analysis was performed by ANOVA followed by Tukey post-hoc test (\**P* < 0.05; \*\**P* < 0.01; \*\*\**P* < 0.001; N.S., not significant). See also Figure S3.

Importantly, treatment of the lysosomal inhibitor, chloroquine (CQ) (Caporaso et al., 1992; Mielke et al., 1997), almost completely blunted *vha-6* RNAi-induced reduction Aβ aggregations in GMC101 worms (Figure S3B), confirming a lysosome-dependent regulatory mechanism. The prototypical aging-associated decline in movement and paralysis of GMC101 worms was also fully normalized by *vha-6* RNAi in an ELT-2-dependent manner (Figure 5D).

Likewise, *vha-6* RNAi reduced the aging-associated formation of polyQ and mutant superoxide dismutase 1 (SOD1) aggregates in *C. elegans* models of HD and Amyotrophic Lateral Sclerosis (ALS) (Morley et al., 2002; Gidalevitz et al., 2009), respectively (Figures 5E, 5F, S3C, and S3D). Moreover, the overall fitness, evaluated by alleviated paralysis and increased movement, was also improved by *vha-6* RNAi in the polyQ or ALS animal models (Figures 5G and 5H).

Furthermore, in a pan-neuronal human Aβ1-42 expressing worm strain GRU102 which displays age-dependent neuromuscular behavior impairments reminiscent of AD pathogenesis (Fong et al., 2016), *vha-6* RNAi improved the intermediate-term memory (Kauffman et al., 2010; Stein and Murphy, 2014) of GRU102 worms to a level similar to that in the non-Aβ expressing control worms (Figure 5I). Interestingly, *vha-6* RNAi even had a modest beneficial effect on memory in control animals (Figure 5I).

Finally, in line with previous studies (Budovskaya et al., 2008; Mann et al., 2016; Sun et al., 2020), the transcripts of the lysosomal protease genes, including *cpr-5*, *cpr-8*, *ctsa-1* and *asp-10*, as well as *vha-6* and *elt-2*, were progressively down-regulated with age in worms fed with control RNAi (Figure 5J). In contrast, in the long-lived *vha-6* RNAi worms, the expression levels of these LySR- related transcripts were increased and sustained throughout the life history; the fitness/movement was also overall improved, which were entirely blunted by *elt-2* RNAi (Figures 5J and 5K). Collectively, these results suggest that activation of LySR by *vha-6* RNAi reduces protein aggregates and extends the healthspan of *C. elegans,* a process highly dependent on ELT-2.

### Overexpression of GATA4/GATA6 reduces Aβ proteotoxicity in mammalian cells

Next, we extended our investigation to a human neuroblastoma SH-SY5Y cell line expressing the Swedish K670N/M671L mutation of the amyloid precursor protein (APPSwe) (Zheng et al., 2011). Infection of the SH-SY5Y APPSwe cells with lentiviral particles expressing full-length human GATA4 or GATA6, the mammalian orthologs of ELT-2 (Shapira et al., 2006; Lentjes et al., 2016), resulted in a remarkable reduction of intracellular Aβ deposits as revealed with an Aβ17-24 specific (4G8) antibody (Figure 6A). This improvement was accompanied by increased protein levels of Cathepsin B and D (CTSB and CTSD) (Figure 6B), and was fully blunted by the lysosomal inhibitor CQ (Figure S4A) (Caporaso et al., 1992; Mielke et al., 1997). Also, the mRNA levels of 8 (i.e., Cathepsin A, D, E, B, F, O, K and V) out of 15 all known lysosomal cathepsins (Cathepsin G and W were not detectable in SH-SY5Y cells) (Figure 6C) (Yadati et al., 2020), covering all the three types of serine, aspartic, and cysteine protease, were increased upon GATA4/GATA6 overexpression. Of note, the mRNA level of APPSwe was not affected (Figure 6C). Furthermore, increased CTSB activity was detected in the total cell lysates of SH-SY5Y APPSwe cells with enforced expression of GATA4 or GATA6 (Figure 6D).

**Figure 6.**
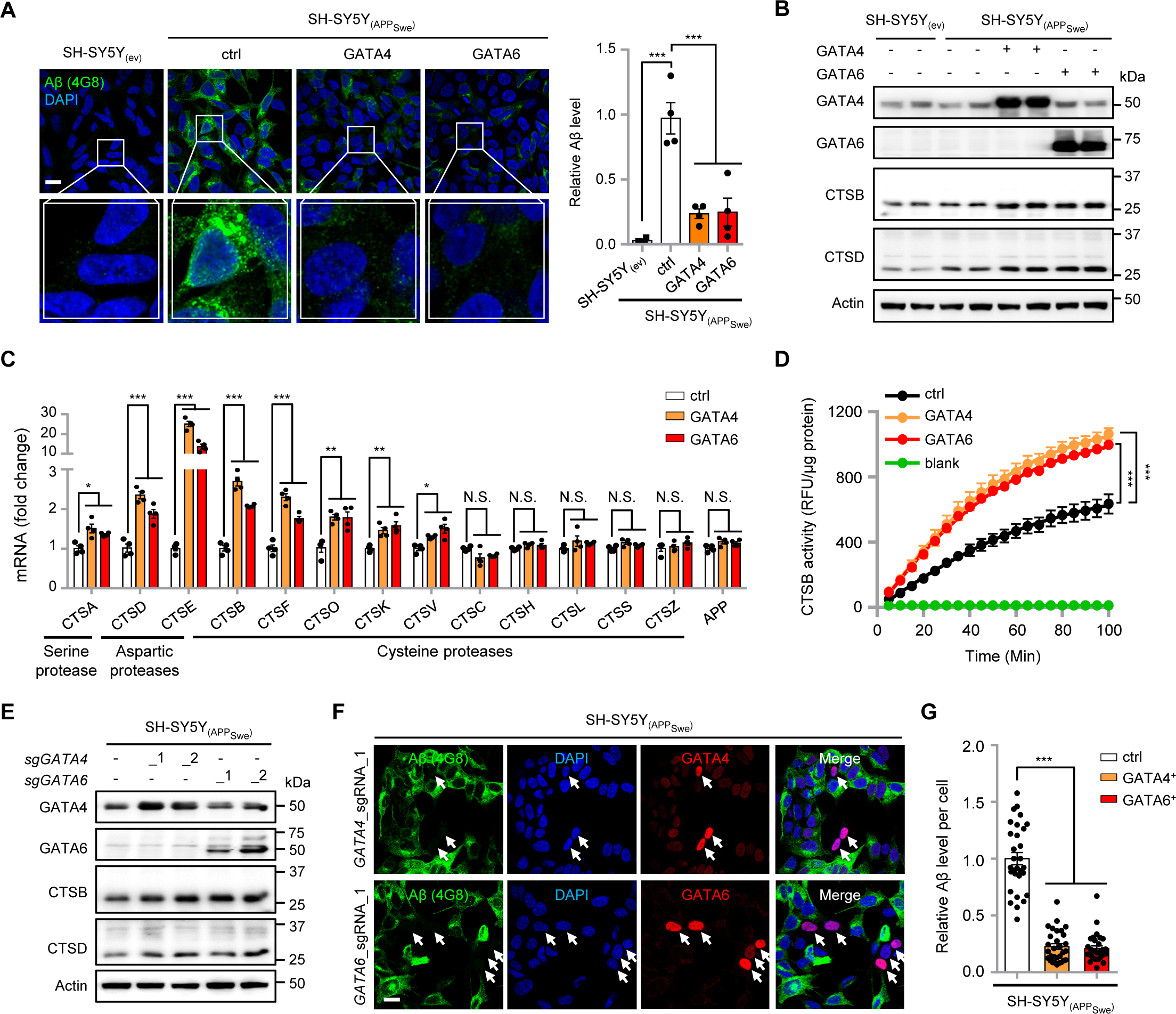
Overexpression of GATA4/GATA6 is sufficient to increase the expression of multiple lysosomal proteases and reduce amyloid-β proteotoxicity in mammalian cells. (A) Images and statistical results (*n* = 4 independent experiments) showing the relative Aβ level of the SH-SY5Y APP_Swe_ cells overexpressed with control (ctrl), GATA4 or GATA6. Scale bar, 20 μm. (B and C) Western blots (B) and qRT-PCR (*n* = 4 biologically independent samples) (C) results of SH-SY5Y APP_Swe_ cells overexpressed with control (ctrl), GATA4 or GATA6. (D) Increased activity of Cathepsin B (*n* = 3 independent experiments) in the whole-cell lysates of SH-SY5Y APP_Swe_ cells overexpressed with GATA4 and GATA6, as compared to that in control cells. RFU, relative fluorescence units. Lysates were replaced by the assay buffer in blank conditions. (E to G) Western blots (E) and immunostaining analysis (F) of Aβ and GATA4 or GATA6 levels in SH-SY5Y APP_Swe_ transfected with sgRNA for endogenous overexpression of GATA4 or GATA6. Note that in cells with high expression of GATA4 or GATA6 (arrows, GATA4/GATA6 positive), few Aβ aggregates were detected as compared to the neighboring cells with basal expression of GATA4 or GATA6, as statistically summarized in (G) (*n* = 30 cells for each group). Scale bar, 20 μm. Error bars denote SEM. Statistical analysis was performed by ANOVA followed by Tukey post-hoc test (\**P* < 0.05; \*\**P* < 0.01; \*\*\**P* < 0.001, N.S., not significant). See also Figure S4.

As an alternative approach to increase GATA4/GATA6 expression, we took advantage of the dCas9-CRISPRa transcriptional activation system (Konermann et al., 2015), and designed two independent single-guide RNAs (sgRNAs) for endogenous transcriptional activation of *GATA4* and *GATA6*, respectively (Figures 6E and S4B). We found that in SH-SY5Y APPSwe cells that were positive for GATA4/GATA6 after transfection with the corresponding sgRNAs, only minor Aβ deposits were detected as compared to their neighboring cells with basal expression of GATA4/GATA6 (Figures 6F, S4C and S4D). Finally, by chromatin immunoprecipitating endogenous GATA4, coupled with quantitative PCR in SH-SY5Y cells, we detected a direct binding of GATA4 to the promotors of lysosomal cathepsins (e.g., Cathepsin A, D, E, B and F) that were responsive to GATA4/GATA6 overexpression (Figure S4E). Together, these results suggest that enforced expression of GATA4/GATA6 is sufficient to induce the expression of multiple lysosomal proteases and reduce amyloid-β proteotoxicity in mammalian cells.

## DISCUSSION

Extensive work has described and characterized key transcriptional responses to promote the homeostasis of mitochondria and endoplasmic reticulum (ER) and regulate aging in multiple organisms (Zhao et al., 2002; Kapulkin et al., 2005; Durieux et al., 2011; Walter and Ron, 2011; Shpilka and Haynes, 2018). In contrast, little was known about the responses or pathways to surveil and improve the function of lysosomes to counteract aging and aging-associated diseases (Lakpa et al., 2021). Here, we discovered a previously uncharacterized longevity-linked lysosomal surveillance response (LySR) that can be activated by RNAi of specific v-ATPase subunits (e.g., *vha-6* RNAi) in *C. elegans*. Typified by the induction of a large panel of lysosome/proteolysis-related genes (e.g., *cpr-5* and *cpr-8*) and almost exclusively regulated by the GATA transcription factor ELT-2, LySR activation improves proteostasis, reduces protein aggregates and extends the health as well as lifespan in several *C. elegans* models of neurodegenerative diseases and of normal aging. In mammalian cells, a similar beneficial transcriptional response could also be induced by enforced expression of GATA4/GATA6, leading to increased expression of multiple lysosomal proteases (e.g., *CTSB* and *CTSD*) and reduced Aβ proteotoxicity (Figure 7).

**Figure 7.**
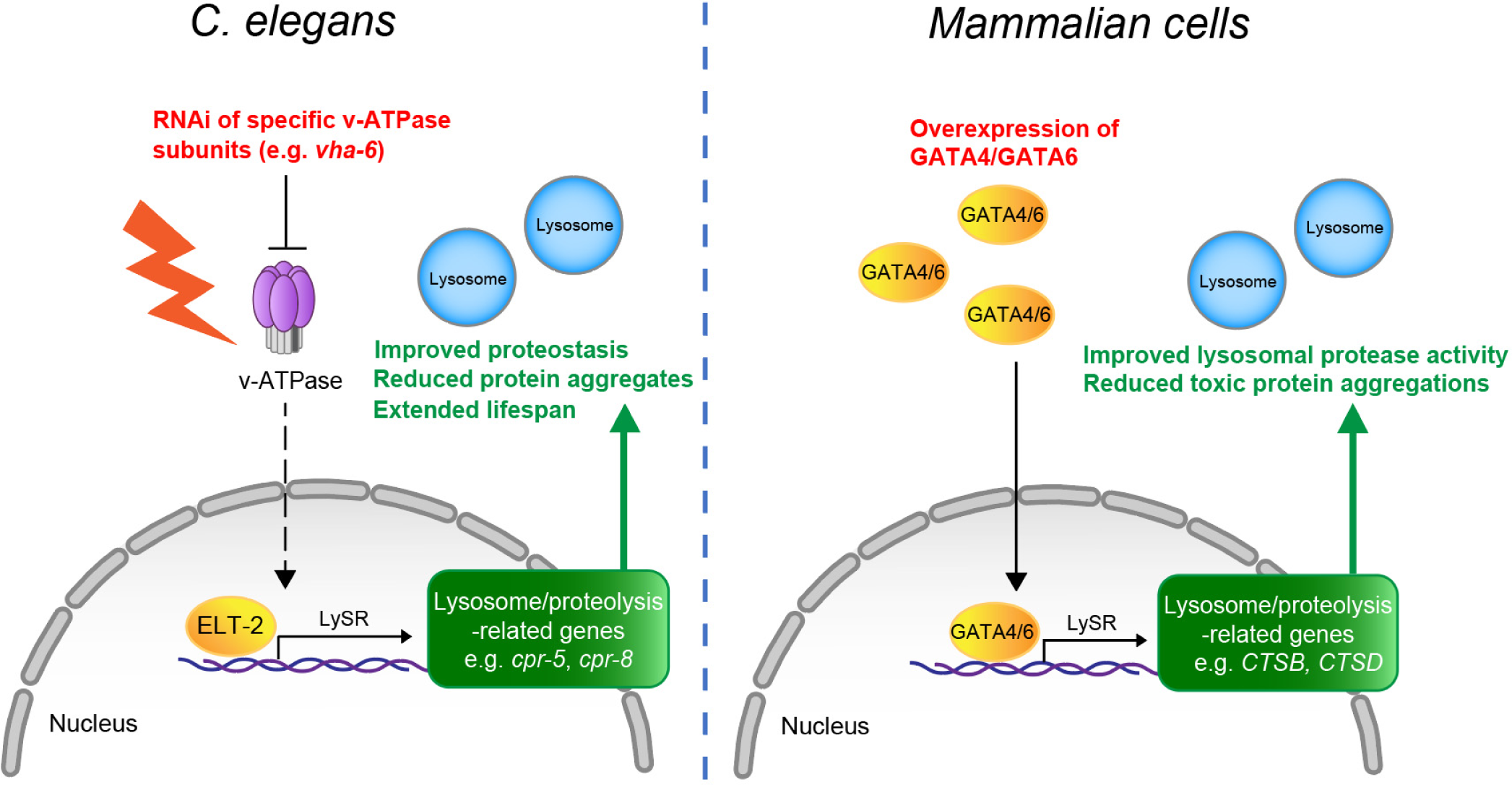
The proposed model for LySR activation and regulation. In *C. elegans*, RNAi of specific lysosomal v-ATPase subunits (e.g., *vha-6* RNAi) activates a beneficial lysosomal surveillance response (LySR) that is controlled by the GATA transcription factor ELT-2. Activation of LySR is typified by the up-regulation of a large panel of lysosome/proteolysis-related genes (e.g., *cpr-5* and *cpr-8*) and thereby improves proteostasis, reduces protein aggregates and extends the lifespan/healthspan in normal aging worms and in several worm models of neurodegenerative diseases. In mammalian cells, a similar beneficial transcriptional response could also be induced by overexpression of GATA4 or GATA6, leading to increased expression of multiple lysosomal proteases (e.g., *CTSB* and *CTSD*) and reduced toxic protein aggregations.

How the RNAi of different subunits of v-ATPase leads to distinct lifespan and gene expression changes in *C. elegans* remains an important direction for future work. For example, *vha-6*, *vha-16* and *vha-19* all encode one of the subunits of the v0 domain of v-ATPase, but only *vha-6* RNAi extends animal lifespan (Figures 1A, 1F and 1G). The difference in lifespan is unlikely due to the insufficiency of some of the RNAi (e.g., *vha-6*, *vha-16* or *vha-19* RNAi) used, for three reasons: first, similar overall number of the up- or down-regulated transcripts were found in worms fed with *vha-6*, *vha-16* or *vha-19* RNAi and the majority (∼70%) of them were actually overlapping (Figures 1I and S1O), indicating that the lysosomes are consistently stressed; second, we have used four doses and three different RNAi of *vha-6*, and the lifespan extension effect was reliably reproduced (Figures 2C and 2F); third, compared to *vha-6* RNAi fed worms, the worms fed with *vha-16* or *vha-19* RNAi were even a bit smaller in size and showed more induction of autophagy-related transcripts (Figure 1K and 1L), suggesting that the LySR is rather specifically activated upon RNAi of certain, e.g., *vha-6*, *vha-8*, *vha-14*, *vha-15*, *vha-20* (Figures 1L, 2A and 3G), but not all v-ATPase subunits. Another possible explanation is that different v-ATPase subunits are expressed at different basal levels in different tissues, and that some subunits may be partially redundant with others (Lee et al., 2010). Indeed, based on the GFP-reporter expression patterns of v-ATPase subunits, most v-ATPase subunits have distinct tissue-specific expression in *C. elegans*; some are primarily expressed in the H-shaped excretory cell (e.g., *vha-1*, *vha-2, vha-4*, *vha-5*, *vha- 11*, *vha-17*) (Oka et al., 1997; Oka and Futai, 2000; Oka et al., 2001; Lee et al., 2010); *vha-6* is almost exclusively expressed in the intestine (Oka et al., 2001; Allman et al., 2009); *vha-7* is enriched in the hypodermis, uterus and spermatheca (Oka et al., 2001); *vha-8* is highly expressed in the hypodermis, intestine and excretory cells (Choi et al., 2003; Ji et al., 2006); *vha-15* is expressed in the muscle, intestine and neurons (McKay et al., 2003); *vha-20* in the intestine, excretory cell and amphid neurons (Zima et al., 2015); and *vha-16* and *vha-19* are more-widely present in excretory cell, hypodermis, intestine and pharynx (Hunt-Newbury et al., 2007; Knight et al., 2012).

Despite that ELT-2 is essential for LySR and LySR-associated lifespan extension in response to the RNAi of certain v-ATPase subunits (Figures 3 and 4), overexpression of ELT-2 seems not sufficient to activate the LySR in *C. elegans*. First, in two independent studies (Zhang et al., 2013; Mann et al., 2016), *elt-2* overexpression extended the lifespan of *C. elegans* by only ∼20%, an extent that is much less that of the ∼60% extension of *vha-6*, *vha-8*, *vha-14* or *vha-20* RNAi (Figures 1A-1E). Second, only sporadic induction of LySR genes in ELT-2 overexpression worms was detected at both young and old age, as compared to that in control worms (Figure S2C). These results suggest that either higher ELT-2 expression levels, or additional co-regulators, are required to fully activate LySR at organismal level in *C. elegans*. In support of the model that other co- factors may cooperate with ELT-2 for LySR activation, while ELT-2 constitutively co-localized with the nuclear DAPI signal under both basal and *vha-6* RNAi conditions (Figure 4F), more ELT- 2 was found to bind to the promoters of LySR-associated lysosomal proteases during LySR activation (Figure 4E). Meanwhile, the DAPI signal appeared more scattered in response to *vha-6* RNAi (Figures 4F and 4G), suggesting that some post-modifications of ELT-2 or histone/DNA epigenetic modification changes may play a role in facilitating the binding of ELT-2 to the LySR gene promoters for ultimate LySR activation. Admittedly, other transcriptional factors, such as DPY-27 and SKN-1 (Figure 3A) (Chuang et al., 1994; Blackwell et al., 2015), may also contribute to the induction of LySR genes in *C. elegans*.

Interestingly, in the human neuroblastoma SH-SY5Y APPSwe cells, we found that overexpression of the ELT-2 mammalian orthologs, GATA4 and GATA6, is sufficient to induce the expression of multiple lysosomal cathepsins including CTSB (the mammalian ortholog of worm CPR-5 and CPR-8) and CTSD, at both mRNA and protein levels (Figures 6B and 6C). Notably, GATA4/GATA6 overexpression remarkably increases the lysosomal protease CTSB activity, and alleviates Aβ proteotoxicity (a process furthermore completely blunted by the lysosomal inhibitor CQ) (Figures 6A, 6F, S4B, S4A and S4C), phenocopying the beneficial effects of *vha-6* RNAi in *C. elegans* (Figures 5C, 5E, 5F, and S3B-S3D). The different impacts of ELT-2 and GATA4/GATA6 overexpression thus suggest a conserved, yet might mechanistically different, regulatory function of GATA transcription factors in LySR activation across species from worms to humans.

Similar to the UPR^mt^ induced by *cco-1* RNAi (Durieux et al., 2011), the LySR activated by *vha-6* RNAi is also likely cell-non-autonomous in *C. elegans*, especially considering that *vha-6* is predominantly expressed in the intestine while the aggregation-prone proteins in the AD, HD and ALS worm models applied in the current study were expressed either under the muscle (*unc-54*)- or neuro (*unc-119*)- specific promoter (Morley et al., 2002; Gidalevitz et al., 2009; McColl et al., 2012; Fong et al., 2016). Thus, it is highly possible that systematic activation of the LySR may furthermore require tight coordination of multiple tissues, which at least includes the intestine, muscle and neurons. In agreement with such a hypothesis, many cathepsins have been shown to be secreted into the extracellular space and serum, and thus implicated in a wide range of physiological processes in mammalian systems via tissue-to-tissue crosstalks (Vidak et al., 2019; Yadati et al., 2020). Likewise, a recent study suggested that induced lysosomal lipolysis in peripheral fat storage tissue activated a neuropeptide signaling pathway in the nervous system to promote longevity (Savini et al., 2022). Interestingly, unlike *cco-1*, whose knockdown induces UPR^mt^ and extends the worm lifespan only when the RNAi was fed during larval development (Durieux et al., 2011), *vha-6* RNAi extends *C. elegans* lifespan even when the knockdown starts at the L4/young adult stage (Figure S1R) (Curran and Ruvkun, 2007), indicating that the anti-aging effects of LySR are likely maintained even in adult or aged animals.

Collectively, by employing genetic, molecular biology and bioinformatic approaches applied in normal aging and multiple neurodegenerative disease models of *C. elegans* as well as in mammalian cells, we here delineated a key mechanism that boosts lysosomal function, promotes protein homeostasis and extends animal healthspan. Cathepsin B, one of the crucial enzymes involved in the degradation of neurotoxic proteins in AD, HD and ALS mouse models (Kikuchi et al., 2003; Liang et al., 2011; Embury et al., 2017), belongs to the LySR network and is induced by overexpression of GATA4/GATA6 in mammalian cells. Targeting the LySR to improve lysosomal homeostasis and reduce proteotoxicity may therefore potentially provide protection against normal aging and neurodegenerative diseases.

## Methods

### C. elegans strains

The N2 (Bristol) strain was employed as the wild-type strain. IA123 (ijIs10[cpr- 5::GFP::lacZ+unc-76(+)]), CB1370 [daf-2(e1370)], CF1038 [daf-16(mu86)], VC222 [raga-1(ok386)], RB754 [aak-2(ok524)], DA465 [eat-2(ad465)], VC3201 [atfs-1(gk3094)], OP56 (gaEx290 [elt-2::TY1::EGFP::3xFLAG(92C12) + unc-119(+)]), CL2122 (dvIs15 [pPD30.38] unc-54(vector) + (pCL26) mtl-2::GFP]), GMC101 (dvIs100 [unc-54p::A-beta-1-42::unc-54 3′- UTR + mtl-2p::GFP]), AM140 (rmIs132 [unc-54p::Q35::YFP]), AM725 (rmIs290 [unc-54p::Hsa-sod-1(127X)::YFP]), GRU101(gnaIs1[myo-2p::yfp]), GRU102 (gnaIs1[myo-2p::yfp + unc-119p::Aß1-42]), HZ1683 [atg-2(bp576)], HZ1684 [atg-3(bp412)], HZ1687 [atg-9(bp564)] and HZ1688 [atg-13(bp414)] were provided by the Caenorhabditis Genetics Center (CGC, University of Minnesota). All worm strains were routinely maintained at 20 °C (except for the CB1370 [daf-2(e1370)] strain, which was maintain at 15 °C) on Nematode Growth Medium (NGM) or high growth medium [HGM: NGM recipe modified as follows: 20 g/L Bacto-peptone, 30 g/L Bacto-agar, and 4 mL/L cholesterol (5 mg/mL in ethanol); all other components same as NGM)] plates, with Escherichia coli (E. coli) OP50 as the food source (Brenner, 1974).

### RNA interference, imaging and compound treatment of *C. elegans*

For RNAi experiments, worms were fed with *E. coli* strains HT115(DE3) containing an empty vector L4440 or expressing double-strand RNAi. RNAi clones were obtained from either Ahringer or Vidal library and verified by sequencing or qRT-PCR before use. The *vha-6* RNAi clone from Vidal library (11038-D9, *vha-6* RNAi_1) was used for all experiments related to *vha-6* RNAi unless otherwise indicated. The other two *vha-6* RNAi clones used were both from the Ahringer library with the accession codes: II-7F06 for *vha-6* RNAi_2, and II-7F04 for *vha-6* RNAi_3.

The RNAi clones for *elt-4*, *elt-6* and *egl-27* were constructed by PCR amplification of cDNAs from total RNA with the following primers: *elt-4*_RNAi_Fw: 5’- TAGATGCTTCTCATCGGAA ACGG-3’, *elt-4*_RNAi_Rv: 5’-CAGTTTCGAAATGCCAGGAGC-3’; *elt-6*_RNAi_Fw: 5’-GAT GCGCTCAGCTTCACAAG-3’, *elt-6*_RNAi_Rv:5’-GAAAACGGCTGCTTGACTGG-3’; *egl- 27*_RNAi_Fw: 5’-ACAAGAACGAGCTGAGCTTGAA-3’, *egl-27*_RNAi_Rv:5’- AAAGACCGTTTGCGTGATGC-3’. The PCR products were then ligated into the L4440 empty vector and transformed into *E. coli* HT115 competent cells.

For RNAi feeding, RNAi bacteria were inoculated and cultured in lysogeny broth (LB) medium with 100 μg/ml ampicillin overnight on a shaker at 37 °C. And then the bacteria were seeded onto RNAi plates (NGM containing 2 mM IPTG and 25 mg/ml carbenicillin) and allowed to form a dry bacterial lawn. Experiments with mixed RNAi was achieved by mixing bacterial cultures, normalized to their optical densities measured at OD600, before seeding. For worm imaging, worms at the last larval stage (Larval stage 4, L4) were picked and transferred onto the RNAi bacteria-seeded plates and incubated at 20 °C to allow overnight egg laying. After 24 h for worm development and egg laying, adult worms were removed from the plates. When the eggs were grown and developed to young adults, 8-10 worms were randomly picked and aligned after placing in a drop of 10 mM tetramisole (Cat. T1512, Sigma) shortly. Fluorescent photos were taken with the same exposure time for each condition using a Nikon SMZ1000 microscope. For aggregate quantification, after the AM140 and AM725 eggs reaching L4 stage, worms were washed off the plate and transferred onto RNAi plate containing 10 μM of 5-FU and allowed to develop to desired age. Worms were randomly picked and imaged after submerging in a drop of 10 mM tetramisole. Aggregates were counted for each worm on day 1, 5 and 8 of adulthood. The GFP intensity of worms was analyzed by using the ImageJ/Fiji 1.53c software.

For chloroquine (CQ) treatment, CQ (Cat. C6628, Sigma) was dissolved in M9 buffer (6 g/L Na2HPO4, 3 g/L KH2PO4, 5 g/L NaCl and 1 mL/L 1M MgSO4 in distilled water) at 400 mM and used as the stock. CQ at final concentration of 1 mM or 5 mM was added to the NGM just before pouring the plates. After RNAi bacteria seeding, synchronized worm eggs obtained by bleaching were then transferred onto the NGM plates and harvested at L4/young adult stage for western blots. In *C. elegans*, some mM levels of CQ are required to functionally inhibit the lysosomal activity, as described previously (Chapin et al., 2015; Zhou et al., 2019).

### DVE-1::GFP and DAPI imaging and quantification

The staining and imaging of 4’,6-diamidino-2-phenylindole (DAPI) in worms was performed as described previously (Porta-de-la-Riva et al., 2012). Briefly, *elt-2p::elt-2::gfp-flag* worms fed on control or *vha-6* RNAi were fixed by ethanol, and stained with DAPI at a final concentration of 2 ng/μl. Worms were then mounted on 2% agarose pads and imaged at 63x using a ZEISS LSM 700 confocal microscope. Quantification of the DAPI signal were performed using ImageJ/Fiji 1.53c software as described (Tian et al., 2016), the image voxels were ranked by DAPI intensity within each nucleus and divided into 4 equal-volume bins. Percentage of total DAPI intensity in each of the bins was then quantified. Analyses were performed in at least 30 nuclei for each condition.

### Lifespan and paralysis analysis

Lifespan assays were conducted as described in the previous study (Houtkooper et al., 2013). Briefly, 5-10 L4 hermaphrodite worms were randomly picked from maintenance plates and transferred onto plates seeded with the indicated RNAi bacteria. After 24 h for worm development and egg laying, adult worms were removed from the plates. Synchronized larvae were raised at 20 °C until they developed into L4 worms. 80-100 L4 worms were randomly picked and transferred onto NGM plates containing 10 μM of 5-FU and seeded with HT115 *E. coli* carrying either empty vector or RNAi clones. Worms were scored every other day and transferred weekly onto freshly bacteria-seeded plates. Those escaped from the plates or had vulva explosion were censored from the assay. All the lifespan assays were performed at least two independent times and similar results were acquired. Paralysis analysis was manually scored after poking, as described previously (McColl et al., 2012), at least 80 total worms were analyzed for each condition.

### RNA extraction and RNA-seq analysis

For worm samples, synchronized worm eggs obtained by bleaching were transferred onto RNAi plates and cultured at for 2.5 days at 20 °C to allow developing to L4/young adult stage. Worms were washed off the plates with M9 buffer for three times and the worm pellets were snap-frozen in liquid nitrogen. To extract total RNA, 1 ml TriPure Isolation Reagent (Cat. 11667165001, Roche) was pipetted to each worm sample. The cell membranes were ruptured by freezing with liquid nitrogen and thawing in a water bath (37 °C) quickly for eight times. And then, total RNAs were extracted using a column-based kit (Cat. 740955.250, Macherey-Nagel). For cells, 1 ml of the TriPure Isolation Reagent (Cat. 11667165001, Roche) was directly added to the cells, and then cell homogenate was transferred to a 1.5 ml Eppendorf tube followed by using the same kit to extract total RNA. RNA-seq was performed by Beijing Genomics Institute (BGI) with the BGISEQ-500 platform. To analyze the RNA-seq results, adaptor sequences, contamination as well as low quality (Phred Score < 20) reads were filtered out from the raw data. Then, qualified reads were mapped to the worm “*Caenorhabditis_elegans.*WBcel235.89” genome with STAR aligner version 2.6.0a and counted by htseq-count version 0.10.0 using the following flags: -f bam -r pos-s no -m union -t exon -I gene_id. Limma-Voom was used to calculate gene differential expressions. Genes with a Benjamini-Hochberg adjusted *P* value < 0.05, and with either log2FC > 1 or < -1 were considered as significantly up- or down-regulated. Genes with significantly up-regulated (adjusted *P* value < 0.05, log2FC > 1) expression in the *vha-6* RNAi condition, and were then down-regulated by more than 25% of the log2FC after *elt-2* RNAi co-treatment, compared with the log2FC of the *vha-6* RNAi condition, were considered as ELT-2-dependent. Functional clustering was performed with the Database for Annotation, Visualization and Integrated Discovery (DAVID) (Huang et al., 2009). Heat-maps were created using Morpheus (https://software.broadinstitute.org/morpheus).

### Binding motif enrichment analysis

The 760 up-regulated genes upon *vha-6* RNAi, but not upon *vha-16* or *vha-19* RNAi, were extracted from the RNA-seq and used as the input dataset. To identify motifs significantly enriched for the promoters of these genes, motif enrichment analysis was performed with HOMER (v4.11) (Duttke et al., 2019), and used the findMotifs.pl script (with start: -2000 bp; end: 2000 bp). The most enriched *de novo* motif of the input genes was compared against a library of known motifs downloaded from the Cis-BP database (catalog of inferred sequence binding preferences) (Weirauch et al., 2014), using PWMEnrich R package (version 4.31.0) (Stojnic and Diez, 2021). The promoters of the input genes were also downloaded from the resource of the HOMER software. Based on these promoter sequences, genomic distribution of the most enriched motif hit with weight score > 6.0 was calculated using pattern matching method of the regulatory sequence analysis tools (RSAT) web server (http://rsat.sb-roscoff.fr/matrix-scan-quick_form.cgi) (Turatsinze et al., 2008).

### Mammalian cell culture

The human SH-SY5Y neuroblastoma cell line (Cat. CRL-2266, ATCC) expressing the APP Swedish K670N/M671L double mutation (APPSwe) was a gift from Dr. Angel Cedazo-Minguez (Karolinska Institute) (Zheng et al., 2011). HEK293T cells (Cat. CRL-3216) were obtained from ATCC and maintained in Dulbecco’s modified Eagle’s medium (DMEM) containing 4.5 g glucose per liter and 10% fetal bovine serum. SH-SY5Y cells were maintained in DMEM/Nutrient Mixture F-12 (DMEM/F-12) containing 4.5 g glucose per liter, 15 mM HEPES, and 10% fetal bovine serum. All cell lines were validated to be free of mycoplasma contamination before use. For lenti- virus-mediated overexpression of GATA4 or GATA6, plasmids expressing GATA4 (Plasmid #120444) and GATA6 (Plasmid #120445) were obtained from addgene. Lenti-virus expressing GATA4 or GATA6 was packed with the 2^nd^ generation lentiviral packaging plasmids, psPAX2 (Plasmid #12260, addgene) and pMD2.G (Plasmid #12259, addgene) with Lipofectamine 2000 (Cat. 11668019, ThermoFisher) in HEK293T cells. Virus supernatant was then harvested 48 h post-transfection, filtered with a 0.45 μm PVDF filter (Cat. SLHVM33RS, Merck) and stored at - 80 °C until use. For dCas9-CRISPRa-mediated GATA4 or GATA6 overexpression, sgRNAs for human GATA4 (sgRNA_1, 5’-GCGCAGGCTGCGGGACTGTG-3’; sgRNA_2, 5’- GCCCAGCGGAGGTGTAGCCG-3’) and GATA6 (sgRNA_1, 5’- GCCGAAGGGTGCCAGGCTGT-3’; sgRNA_2, 5’-AGGGCTCGGTGAGTCCAATC-3’) were cloned to the sgRNA(MS2) cloning backbone plasmid (Plasmid #61427, addgene), followed by co-transfection with plasmids expressing MS2-P65-HSF1 (Plasmid #61426, addgene) and dCAS- VP64 (Plasmid #61425, addgene), as described previously (Konermann et al., 2015). Transfection of SH-SY5Y cells was conducted with the Lipofectamine 3000 Transfection Reagent (Cat. L3000015, ThermoFisher). SH-SY5Y cells infected with lenti-virus particles expressing GATA4/GATA6, or transfected with sgRNAs for GATA4/GATA6, were analyzed for RT-qPCR, western blots or imaging 72 h after viral infection or transfection, no antibiotic selection was applied throughout the experiments.

### Protein extraction and western blots

Proteins were extracted with Radio-immunoprecipitation assay (RIPA) buffer containing protease and phosphatase inhibitors as previously described (Houtkooper et al., 2013). Western blots were carried out with antibodies against green fluorescent protein (GFP) (Cat. 2956, CST, 1:1,000, RRID:AB_1196615), β-amyloid 1–16 (6E10) (Cat. 803001, BioLegend, 1:1,000, RRID:AB_2564653), Tubulin (Cat. T5168, Sigma, 1:2,000, RRID:AB_477579), GATA-4 (D3A3M) (Cat. 36966, CST, 1:1,000, RRID:AB_2799108), GATA-6 (D61E4) (Cat. 5851S, CST, 1:1,000, RRID:AB_10705521), CTSB (Cat. 31718, CST, 1:1,000, RRID:AB_2687580), CTSD (Cat. 2284, CST, 1:1,000, RRID:AB_10694258), ACTIN (Cat. A5441, Sigma, 1:2000, RRID:AB_476744). Horseradish peroxidase (HRP)-labeled anti-rabbit (Cat. 7074; CST, 1:5,000, RRID:AB_2099233) and anti-mouse (Cat. 7076; CST; 1:5,000, RRID:AB_330924) secondary antibodies.

### Imaging of SH-SY5Y cells

SH-SY5Y cells were seeded on glass cover slips one day before experiment and treated with or without 50 μM chloroquine (CQ, Cat. C6628, Sigma) for 24 h. Cells were then fixed with 4% of formaldehyde, permeabilized with 0.2% Triton X-100, blocked with 5% BSA, and stained with antibodies to β-amyloid 17-24 (4G8) (Cat. 800709, BioLegend, 1:400, RRID:AB_2565325), GATA-4 (D3A3M) (Cat. 36966, CST, 1:250, RRID:AB_2799108), GATA-6 (D61E4) (Cat. 5851, CST, 1:250, RRID:AB_10705521), lysosome marker LAMP1 (Cat. 328626, Biolegend, 1:500, RRID:AB_11203537) or DAPI. The secondary antibodies used were Goat anti-Mouse Alexa Fluor™ 488 (Cat. A-11029, ThermoFisher, RRID:AB_2534088) and Donkey anti-Rabbit Alexa Fluor™ 568 (Cat. A-10042, ThermoFisher, RRID:AB_2534017). Images with the same parameter settings for each condition were then acquired at 63x using a ZEISS LSM 700 confocal microscope.

### Cathepsin B activity measurement

The enzymatic activity of Cathepsin B was measured by using a Cathepsin B Activity Assay Kit (Fluorometric) (Cat. ab65300, abcam). Briefly, SH-SY5Y cells were washed once with cold PBS and resuspended in a Cathepsin B Cell Lysis Buffer within the kit. After incubation on ice for 30 min, the lysates were centrifuged for 5 min at 4°C to remove any insoluble materials. Protein concentration was then determined by using a DC™ Protein Assay Reagent kit (Cat. 5000122, Bio-RAD). Samples were then normalized with the Cell Lysis Buffer based on the protein concentration. Equal volumes of samples were subsequently mixed with the Cathepsin B Reaction Buffer, followed with the addition of the substrate Ac-RR-AFC (200 µM final concentration), the fluorescence units at Ex/Em = 400/505 nm were then monitored every 5 min for a total of 100 min at 37°C and protected from light.

### Quantitative RT-PCR and ChIP-qPCR

Worms were collected and the total RNA was extracted with the same method mentioned above as for RNA-seq. 1000 ng of RNA was used for cDNA synthesis using the Reverse Transcription Kit (Cat. 205314, Qiagen). Quantitative real-time PCR (qRT-PCR) was performed with the LightCycler 480 SYBR Green I Master kit (Cat. 04887352001, Roche). Primers for *pmp-3* and *ACTIN* were used as housekeeping for *C. elegans* and human cells, respectively. ChIP-qPCR was carried out as previously described (Li et al., 2021). Briefly, *elt-2::TY1::EGFP::3xFLAG* worms were fixed with 1% formaldehyde solution for 15 min, and quenched by glycine. After a total 15 min sonication, immunoprecipitations were performed using the anti-FLAG M2 beads (Cat. A2220, Sigma) in RIPA buffer. ChIP for human endogenous GATA4 in SH-SY5Y cells was carried out using the GATA-4 (G-4) (Cat. sc-25310, Santa Cruz, 1:50, RRID:AB_627667). All primers for qRT-PCR or ChIP-qPCR are as indicated in Table S3.

### Thrashing/movement analysis

Worms were randomly picked from the culture plates and ∼15 worms were used for the thrashing assay of each condition. In brief, a single worm was placed in a drop of M9 buffer on a glass slide and allowed to acclimatize to the environment for 30 seconds. One movement of the worm that swung its head to the same side was considered as one thrash, and the frequency of thrashes was counted for 30 seconds as previously described (Brenner, 1974; Currey and Liachko, 2021).

### Positive olfactory associative memory assays

Vector control or *vha-6* RNAi treated wild-type GRU101 and GRU102 animals were trained and tested for intermediate-term memory at Day 4 of adulthood as previously described (Kauffman et al., 2010; Stein and Murphy, 2014). Briefly, synchronized Day 4 adult hermaphrodites were washed from HGM RNAi plates with M9 buffer, allowed to settle by gravity, and washed again with M9 buffer. After washing, the animals were starved for 1 h in M9 buffer. For intermediate-term memory training, worms were then transferred to 10 cm NGM conditioning plates (seeded with OP50 *E. coli* bacteria and with 12 μl 10% 2-butanone (Acros Organics) in ethanol streaked across the lid with a pipette tip for 1 h. After conditioning, the trained population of worms were transferred to 10 cm NGM plates with fresh OP50 bacteria for a 60 min interval before testing worms for intermediate-term memory performance by chemotaxis to 10% butanone previously described chemotaxis assay conditions (Bargmann et al., 1993). Chemotaxis indices were calculated as follows: **(#worms_Butanone_ - #worms_Ethanol_)/(Total #worms)**. Performance index is the change in chemotaxis index following training relative to the naïve (untrained) chemotaxis index which was determined using a subpopulation of animals. The calculation for Performance Index is: **Chemotaxis Index_Trained_ -Chemotaxis Index_Naive_**.

### Statistical analysis

Sample sizes were chosen based on studies with similar experimental design and on the known variability of the assay. All experiments, except for the RNA-seq, were repeated at least twice and similar results were acquired. Investigators were not blinded to allocation during experiments and outcome assessment. No samples were excluded, except for the lifespan assays, whereby worms escaped or had vulva explosion were censored. Graphpad Prism 8 software was used to conduct all the statistical analyses. Two-tailed and unpaired Student’s *t*-test was used to determine the differences between two independent groups. For more than two groups of comparisons, analysis of variance (ANOVA) followed by Tukey’s honest significant difference test was performed. One-way ANOVA was used for comparisons between different groups, and two-way ANOVA was used for examining the effect of two independent variables on a dependent variable, for example, age and strain. For survival analyses, Kaplan-Meier method was performed and the significance was calculated using the log-rank (Mantel-Cox) method.

### Data availability

Original materials are available upon request to the corresponding author (J.A.). The raw and processed RNA sequencing data have been uploaded in the NCBI Gene Expression Omnibus (GEO) database with the accession number: GSE196021 (Reviewer token: ijwhiswqpzmbpcf) and GSE196022 (Reviewer token: exqnggqqlxwftuv).

## ACKNOWLEDGMENTS

We thank the Caenorhabditis Genetics Center for providing the *C. elegans* strains. We thank all members of J. Auwerx and K. Schoonjans laboratories for helpful discussions. This work was supported by grants from the EPFL, the European Research Council (ERC-AdG-787702), the Swiss National Science Foundation (SNSF 31003A_179435 and Sinergia CRSII5_202302), and GRL grant of the National Research Foundation of Korea (NRF 2017K1A1A2013124). T.Y.L. was supported by the “Human Frontier Science Program” (LT000731/2018-L). A.W.G. was supported by the United Mitochondrial Disease Foundation (PF-19-0232). X.L. was supported by the China Scholarship Council (201906050019). Y.J.L and Q.W. was supported by the European Molecular Biology Organization (ALTF 1161-2021 and ALTF 111-2021, respectively). R.N.A. was supported by the Whitehall Foundation and is a Glenn Foundation for Medical Research and AFAR Grant for Junior Faculty Awardee. K.M. is supported by a GC-CPEH NIEHS Pilot Project grant (P30-ES030285) awarded to R.N.A.

## AUTHOR CONTRIBUTIONS

T.Y.L., A.W.G., and J.A. conceived the project. T.Y.L. and A.W.G. performed most of the experiments. Y.J.L., Q.W., and T.I.L. contributed to the aggregation study. R.N.A. and K.M. contributed to the behavior analysis of AD worms. T.Y.L., A.W.G., X.L., and A.L. performed data analysis. J.A. supervised the study. T.Y.L., A.W.G., and J.A. wrote the manuscript with comments from all authors.

---

**Figure S1.**
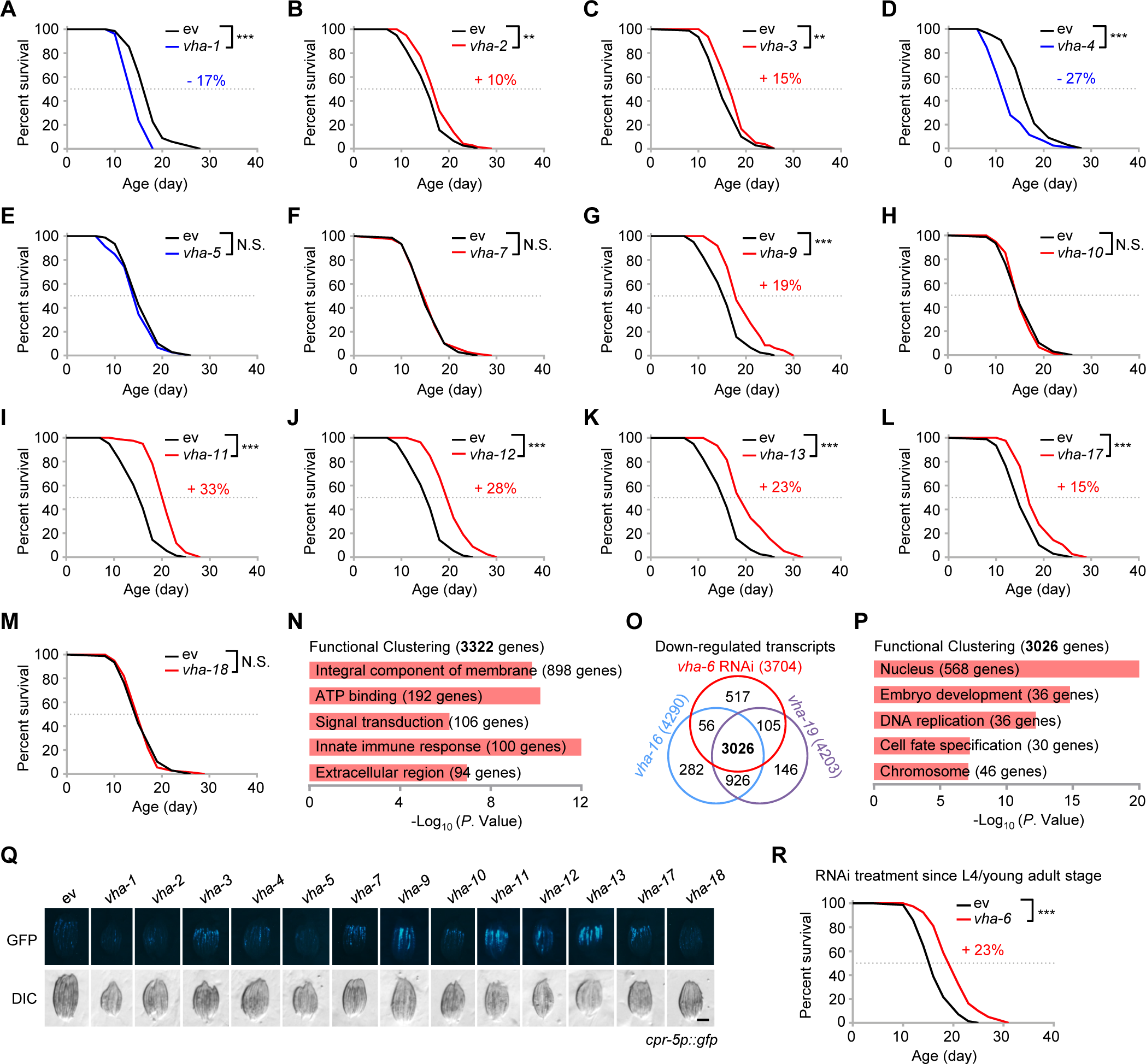
Impact of RNAi targeting different v-ATPase subunits on gene expression and lifespan of *C. elegans*, related to Figure 1. (**A** to **M**) Survival of worms fed with control (ev), or RNAi targeting different v-ATPase subunits. Each v-ATPase RNAi occupied 40%, except for *vha-11* and *vha-12* RNAi, which occupied 10%; control RNAi was used to supply to a final 100% of RNAi for all conditions. The percentages indicate the mean lifespan changes as compared to the control condition. (**N**) Functional clustering of the 3,322 differentially expressed genes (DEGs) that commonly up-regulated in in response to *vha-6*, *vha-16* and *vha-19* RNAi. (**O**) Venn diagram of the down-regulated DEGs in response to *vha-6*, *vha-16* and *vha-19* RNAi. (**P**) Functional clustering of the 3,026 DEGs as indicated in (O). (**Q**) GFP expression levels of *cpr-5p::gfp* worms fed with RNAi targeting different v-ATPase subunits. Scale bar, 0.3 mm. (**R**) *vha-6* RNAi extended wild-type worm lifespan by 23%, even when RNAi treatment started since the L4/young adult stage. Statistical analysis was performed by log-rank test (\*\**P* < 0.01; \*\*\**P* < 0.001; N.S., not significant).

**Figure S2.**
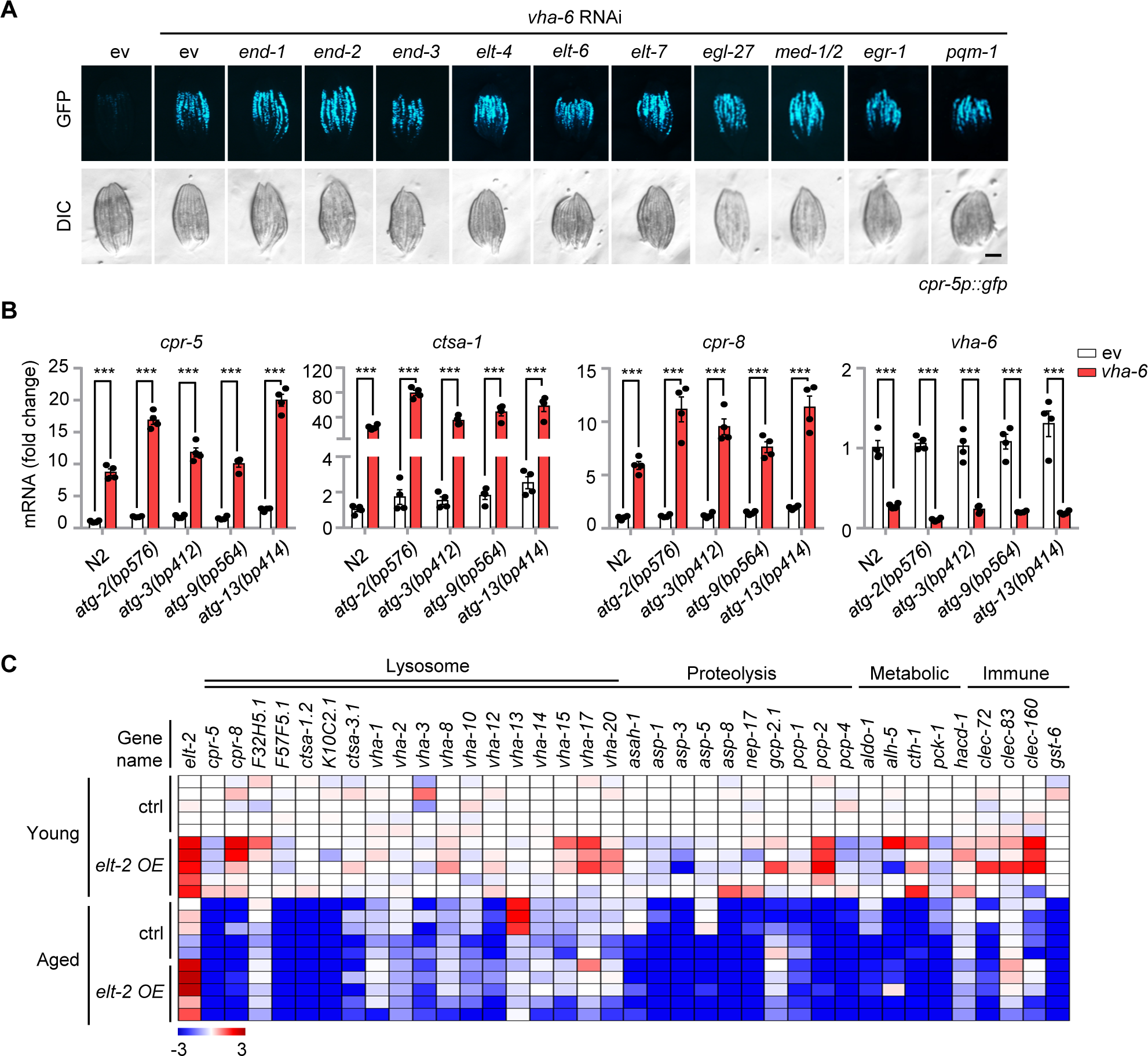
Identification of determinates that regulate LySR activation in *C. elegans*, related to Figure 3. (**A**) RNAi of other GATA transcription factor members, or *pqm-1*, did not affect the GFP expression of *cpr-5p::gfp* worms induced by *vha-6* RNAi. RNAi targeting *vha-6* occupied 40%, RNAi targeting GATA transcription factor members, or *pqm-1*, occupied 60%. Scale bar, 0.3 mm. (**B**) The indicated autophagy defective mutants have a normal induction of representative lysosomal proteases in response to *vha-6* RNAi. qRT-PCR analysis of transcripts (*n* = 4 biologically independent samples) in wild-type (N2) or autophagy defect mutants of *C. elegans* fed with control (ev) or *vha-6* RNAi. (**C**) Heat-map of the relative expression levels of representative LySR genes in control or *elt-2* overexpression (OE) worms at young (L4) or aged (Day 13) stage based on an extant RNA-seq dataset (GSE69263) (Mann et al., 2016). The color represents gene expression differences in log2(fold change, FC) relatively to control RNAi and young (L4) condition. Error bars denote SEM. Statistical analysis was performed by two-tailed unpaired Student’s *t*-test (\*\*\**P* < 0.001).

**Figure S3.**
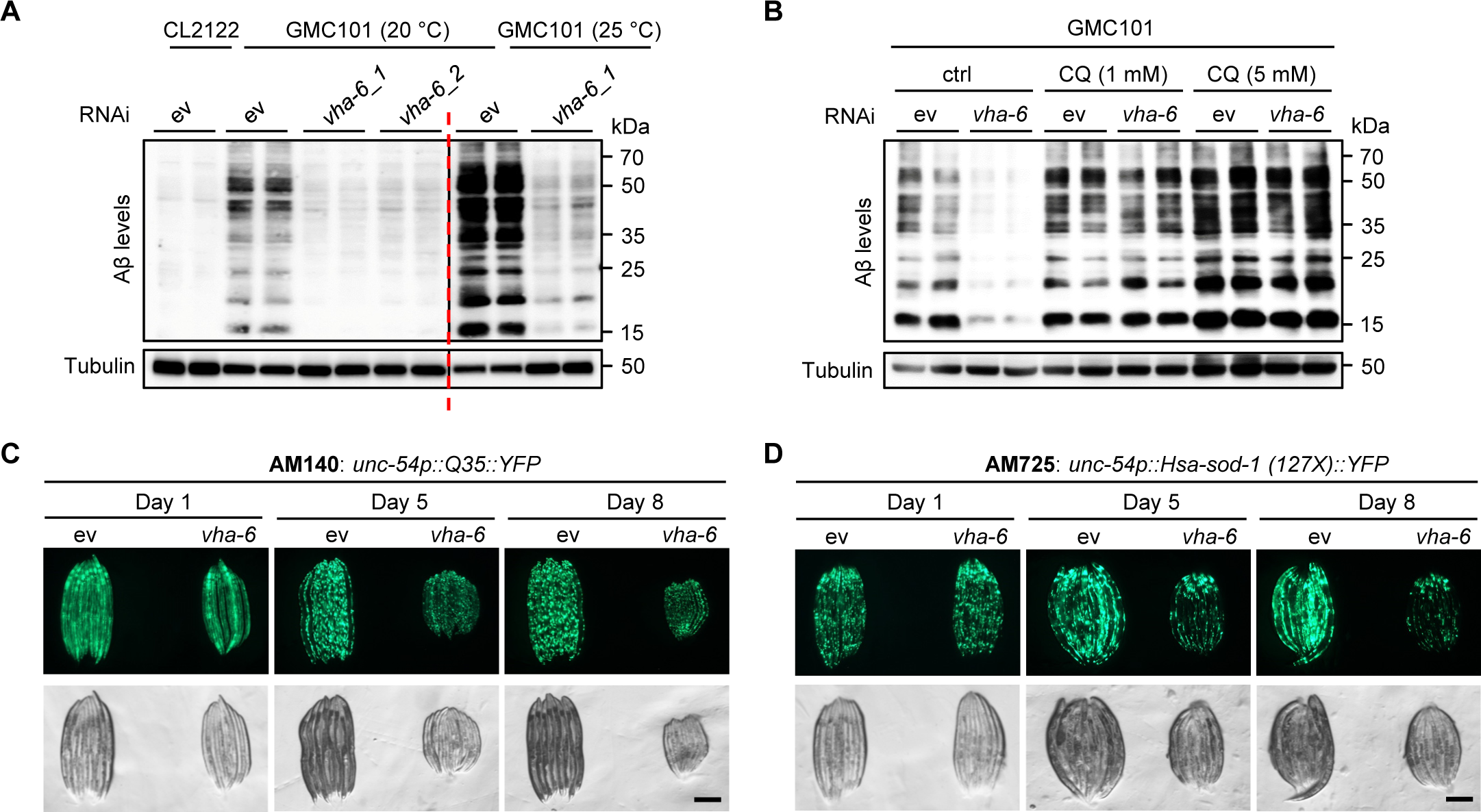
Characterization of worm models expressing different disease-causing protein aggregations with or without *vha-6* RNAi treatment, related to Figure 5. (A) Western blots of CL2122 or GMC101 worms fed with control (ev), *vha-6_1* or *vha-6_2* RNAi, cultured at two different temperatures since L4 stage. (B) Western blots of GMC101 worms fed with control (ev) or *vha-6* RNAi and treated with or without chloroquine (CQ) at a final concentration of 1 mM or 5 mM. (C and D) RNAi of *vha-6* (20%) reduces the aggregate formation in *unc- 54p::Q35::YFP* (polyQ model) (C) and *unc-54p::Hsa-sod-1::YFP* (Amyotrophic Lateral Sclerosis (ALS) model) (D). Scale bars, 0.3 mm.

**Figure S4.**
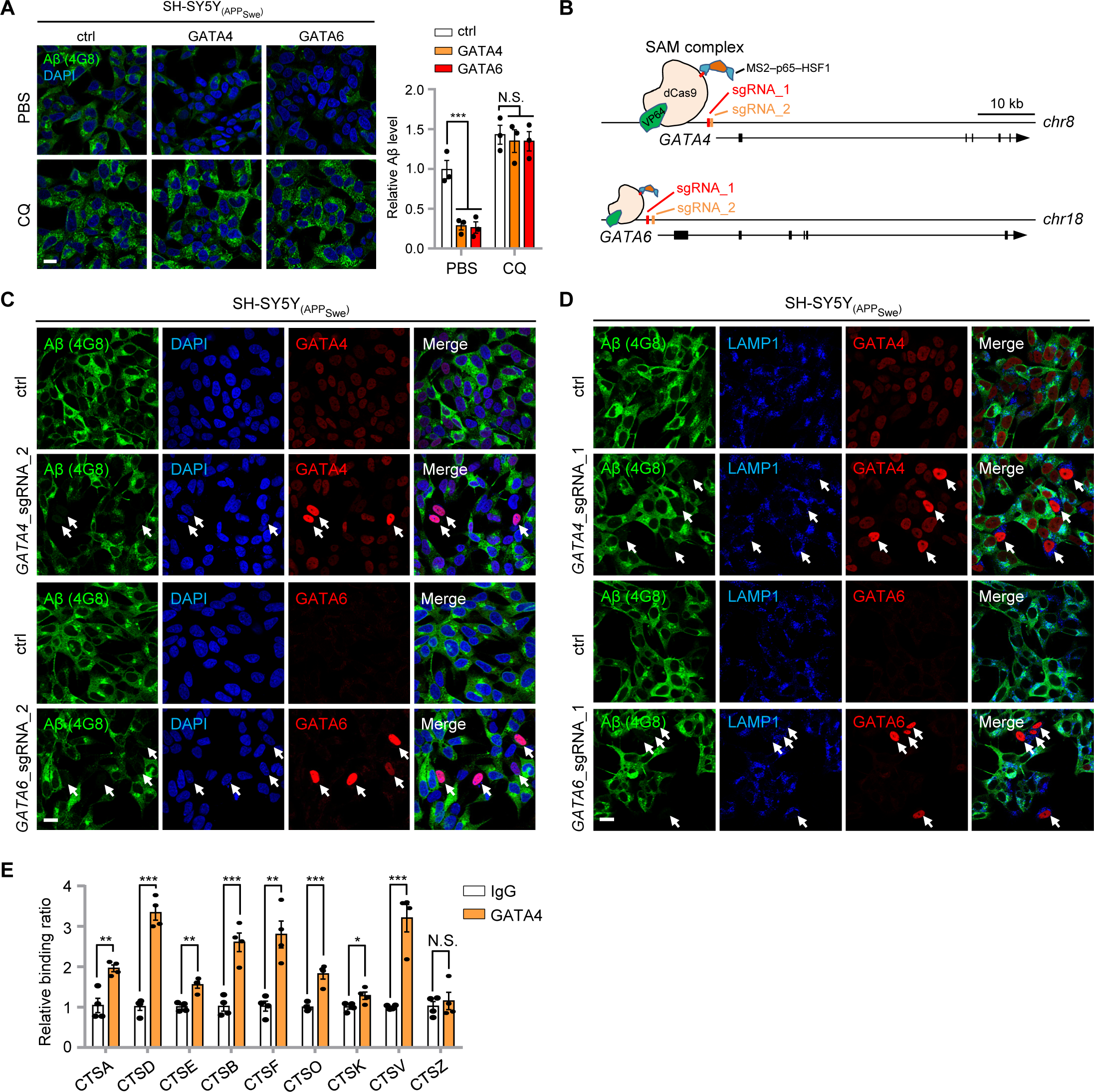
Overexpression of GATA4/GATA6 is sufficient to reduce amyloid-β proteotoxicity in mammalian cells, related to Figure 6. (**A**) Images and statistical results (*n* = 3 independent experiments) showing the relative Aβ level of the SH-SY5Y APP_Swe_ cells overexpressed with control (ctrl), GATA4 or GATA6 and treated with PBS (control) or lysosomal inhibitor chloroquine (CQ) (50 μM) for 24 h. Scale bar, 20 μm. (**B**) Schematic design of the RNA-guided transcription activation of endogenous *GATA4* or *GATA6* using the synergistic activation mediator (SAM) system (Konermann et al., 2015). Two independent CRISPRa sgRNAs upstream of the transcription start site (TSS) of *GATA4* and *GATA6* were designed, respectively. (**C** and **D**) Immunostaining of SH-SY5Y APP_Swe_ transfected with sgRNA for endogenous overexpression of GATA4 or GATA6 co-stained with DAPI (C) or LAMP1 (D). Note that in cells with high expression of GATA4 or GATA6 (arrowed), few Aβ aggregates were detected as compared the neighboring cells with basal expression levels of GATA4 or GATA6. DAPI and LAMP1 stained the nuclei and lysosomes, respectively. Scale bars, 20 μm. (**E**) Direct binding of endogenous GATA4 to the promotors of multiple cathepsin genes. ChIP-qPCR (*n* = 4 biologically independent samples) of SH-SY5Y cells immunoprecipitated with IgG control or anti-GATA4 antibody. Error bars denote SEM. Statistical analysis was performed by ANOVA followed by Tukey post-hoc test (\**P* < 0.05; \*\**P* < 0.01; \*\*\**P* < 0.001; N.S., not significant).

