## Supplemental Table 3 for "A lysosomal surveillance response (LySR) that reduces proteotoxicity and extends healthspan"

**Table S3. List of primers used for qRT-PCR and ChIP-qPCR in this study, related to Methods**

| Species | Application | Gene | Forward primer (5'-3') | Reverse primer (5'-3') |
|---|---|---|---|---|
| *C. elegans* | qRT-PCR | *cpr-5* | TGTGTCGACTCCTGCACTTC | CCATCCGAGGATCTTGACGG |
| *C. elegans* | qRT-PCR | *cpr-8* | TGGCAACAAACAGGGTCAC | ACTCGAGCCCACATTCGTTT |
| *C. elegans* | qRT-PCR | *ctsa-1* | AGCTGCTCCAGCTACGGATA | GTTCCAGGCGTACTCGTTGT |
| *C. elegans* | qRT-PCR | *asp-10* | GTCTCTGCAGAAGTCCACCA | TGGCATCTTTTCCTCCAGCA |
| *C. elegans* | qRT-PCR | *vha-6* | TCCTTTGGAGACTGTCACGC | ACGGCCTTCATGTCAGTTGT |
| *C. elegans* | qRT-PCR | *elt-2* | CCTCTCTGTACGACCCCAGT | ACGCACATCATTCCTCCGTT |
| *C. elegans* | qRT-PCR | *pmp-3* | GTTCCCGTGTTCATCACTCAT | ACACCGTCGAGAAGCTGTAGA |
| *C. elegans* | ChIP-qPCR | *cpr-5* | AGGACTCCGATCAGGCAAAT | GTTCACCGAATAAGCTCCGCT |
| *C. elegans* | ChIP-qPCR | *cpr-8* | AATCCATGAATTTTAGTAGAGCTGC | TTATCCAAAAAGAAATGCAAGCGG |
| *C. elegans* | ChIP-qPCR | *ctsa-1* | AGTGTCGTCAGATGAGACAAAATG | CCTAAAAGGCGGGCTTCACT |
| *C. elegans* | ChIP-qPCR | *asp-10* | CAAGCCGCCCATAAGAGACA | TTCGGCACTATCAGTTCGGC |
| *Human* | qRT-PCR | *CTSA* | TGCTGCTCTCAAAACAAGTGTAA | GGGCCACTTCCTGAAGATT |
| *Human* | qRT-PCR | *CTSD* | GACATCCACTATGGCTCGGG | AGCACGTTGTTGACGGAGAT |
| *Human* | qRT-PCR | *CTSE* | CAGTCCAGCACATACAGCCA | CCACGGTTAGTCCTTCCACAG |
| *Human* | qRT-PCR | *CTSB* | GTGCAGACCGTACTCCATCC | CTCGCTATTGGAGACGCTGT |
| *Human* | qRT-PCR | *CTSF* | CTTCGCGCTGGAGATGTTC | CACGTGTCTTCCGAGCTCAT |
| *Human* | qRT-PCR | *CTSO* | TCCCCAATGTGTCTTTGCCG | CCAGGGGCTTCCCCTTTATT |
| *Human* | qRT-PCR | *CTSK* | GGGGGACATGACCAGTGAAG | CAGAGTCTGGGGCTCTACCT |
| *Human* | qRT-PCR | *CTSV* | GGTACCAGTGGAAGGCAACA | GAAGCCATGTTTCCCTTGGC |
| *Human* | qRT-PCR | *CTSC* | CACCTATCTTGACCTGCTGGG | GCCAGAATTGCCAAGGTCATC |
| *Human* | qRT-PCR | *CTSH* | TGCGCCCAGGACTTCAATAA | CACCATCGCTTCCTCGTCAT |
| *Human* | qRT-PCR | *CTSL* | GCACAGTGGACCAAGTGGAA | AGCTGTGTTTCCCTTCCCTG |
| *Human* | qRT-PCR | *CTSS* | GCCCAGTGTCTGTTGGTGTA | GCCCCAGCTGTTTTTCACAA |
| *Human* | qRT-PCR | *CTSZ* | CGGAGGCATCTATGCCGAAT | CCCATGGTTCACCCCATGAA |
| *Human* | qRT-PCR | *CTSG* | TGAGGCAGGGGAGATCATCG | TGGGTGTTTTCCCGTCTCTG |
| *Human* | qRT-PCR | *CTSW* | TGGGACGCGTTCATAACTGT | GGTACTGCGCAATTCTGTGC |
| *Human* | qRT-PCR | *APP* | GGTTTGGCACTGCTCCTG | CAGTCTGCCACAGAACATGG |
| *Human* | qRT-PCR | *ACTIN* | TCGTGCGTGACATTAAGGAG | GTCAGGCAGCTCGTAGCTCT |
| *Human* | ChIP-qPCR | *CTSA* | GAATCCCTGCCCTAGGTGAATA | CTTGCACAAGCCTCTACATGGT |
| *Human* | ChIP-qPCR | *CTSD* | AGGCTTCCAGAGAGGATGTCT | GGGTGACCCCTGTCCC |
| *Human* | ChIP-qPCR | *CTSE* | AGGACTCCCAAGTTCTCCCC | CCTAACTCTCAGACCTGCCC |
| *Human* | ChIP-qPCR | *CTSB* | GTCGCCACAGCCTGGATAAA | AATCCCAGCTACTCGGGAGA |
| *Human* | ChIP-qPCR | *CTSF* | TTCAGAGATTGCTACGCCGA | CCTGCGCGTTCTCTTGTTT |
| *Human* | ChIP-qPCR | *CTSO* | ACGGAGAAACAGGCTTCAAGA | CACCGGAAGCTGCCTTTTTC |
| *Human* | ChIP-qPCR | *CTSK* | ACTAGGGCTGTGTCCTTCCT | GTATGCAGCATGTGTGCCAG |
| *Human* | ChIP-qPCR | *CTSV* | CAGGAGCCACTGAACGAGAG | GCGCCTACCTCAATACACCC |
| *Human* | ChIP-qPCR | *CTSZ* | CCGGCAAATGAGTCCAGCC | ACTTTGGGCCCCTGCC |
